# Mapping Pesticide-Induced Metabolic Alterations in Human Gut Bacteria

**DOI:** 10.1101/2024.11.15.623895

**Authors:** Li Chen, Hong Yan, Shanshan Di, Chao Guo, Huan Zhang, Shiqi Zhang, Andrew Gold, Yu Wang, Ming Hu, Dayong Wu, Caroline H. Johnson, Xinquan Wang, Jiangjiang Zhu

**Affiliations:** Human Nutrition Program, Department of Human Sciences, The Ohio State University, Columbus, OH 43210, USA; James Comprehensive Cancer Center, The Ohio State University, Columbus, OH 43210, USA; Department of Environmental Health Sciences, Yale School of Public Health, New Haven, CT, USA; State Key Laboratory of Environmental and Biological Analysis, Hong Kong Baptist University, Hong Kong, China; State Key Laboratory for Managing Biotic and Chemical Threats to the Quality and Safety of Agro-products/ Key Laboratory of Detection for Pesticide Residues and Control of Zhejiang, Institute of Agro-product Safety and Nutrition, Zhejiang Academy of Agricultural Sciences, Hangzhou 310021, China; Department of Physiology, Johns Hopkins University School of Medicine, Baltimore, Maryland 21205, USA; Department of Cancer Biology and Genetics, The Ohio State University College of Medicine, Columbus, OH 43210, USA

**Keywords:** Gut bacteria, Pesticide, Metabolomics, Lipidomics, Inflammation

## Abstract

Pesticides can modulate gut microbiota (GM) composition, but their specific effects on GM remain largely elusive. Our study demonstrated that pesticides inhibit or promote growth in various GM species, even at low concentrations, and can accumulate in GM to prolong their presence in the host. Meanwhile, the pesticide induced changes in GM composition are associated with significant alterations in gut bacterial metabolism that reflected by the changes of hundreds of metabolites. We generated a pesticide-GM-metabolites (PMM) network that not only reveals pesticide-sensitive gut bacteria species but also report specific metabolic changes in 306 pesticide-GM pairs (PGPs). Using an *in vivo* mice model, we further demonstrated a PGP’s interactions and verified the inflammation-inducing effects of pesticides on the host through dysregulated lipid metabolism of microbes. Taken together, our findings generate a PMM interactions atlas, and shed light on the molecular level of how pesticides impact host health by modulating GM metabolism.

**Graphical Abstract:** 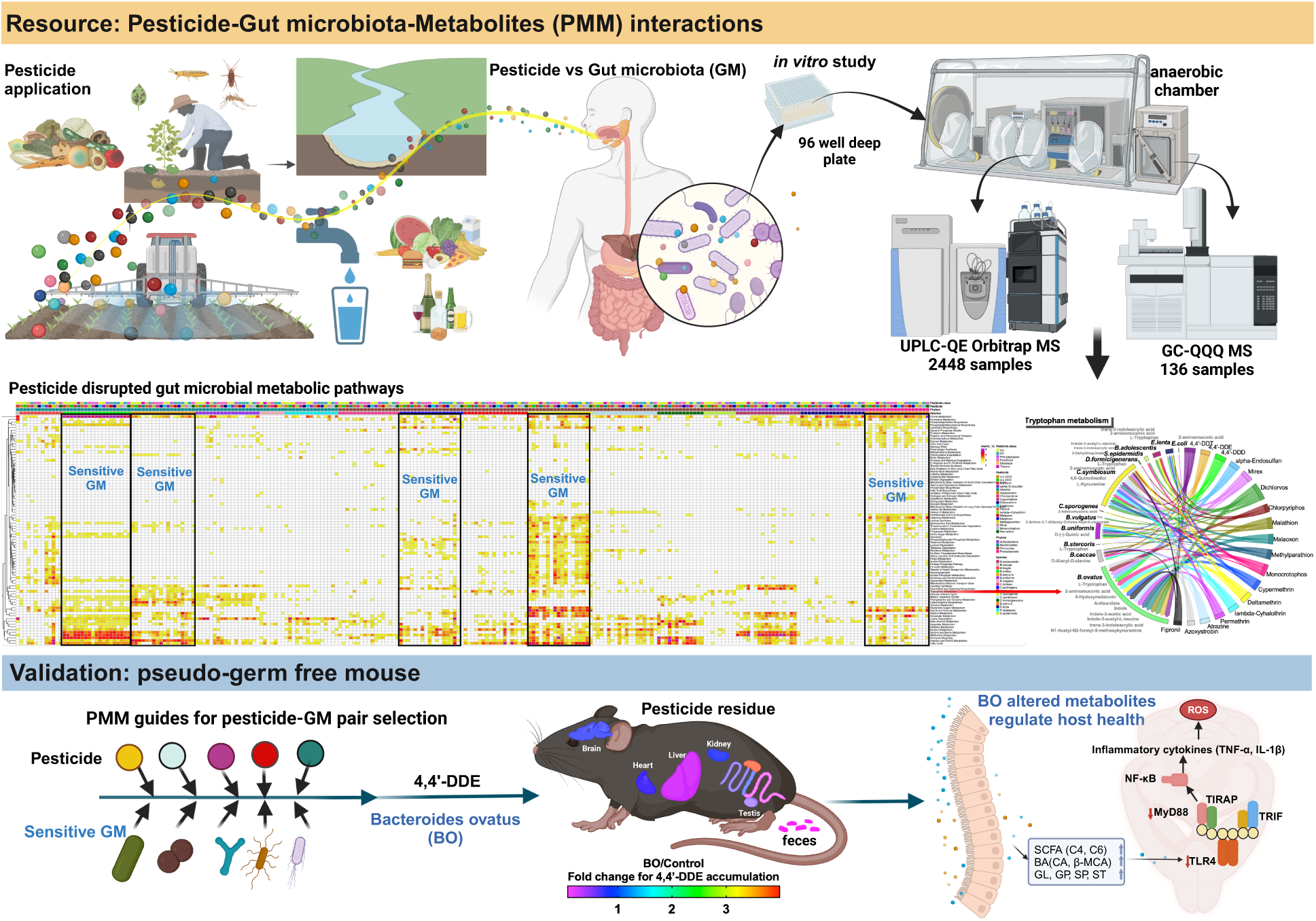

## Introduction

Pesticides are extensively utilized globally to meet the increasing demand for food, and for enhancing the quality of agricultural products. Consequently, concerns regarding the health risks on non-targeted organisms ^1–3^ or humans ^4,5^ associated with pesticides are arising due to their residues in soil, ^6^ water, ^7^ and air.^8^ Human populations often encounter multiple pesticides through their dietary intake or drinking habits.^9,10^ Meanwhile, the gastrointestinal tract (GI), acting as a protective barrier against pathogenic microorganisms and toxins, is also a primary site for pesticides exposure, and plays a crucial role in the metabolic and immune functions of the host.^11,12^ Within the GI, it is well-recognized that a vast gut microbiota (GM) community of 100 trillion microbial cells forms a complex ecosystem,^13^ and this intricate ecosystem maintains a mutualistic relationship with its host. The stability of a healthy GM composition is essential for various physiological processes, including food digestion, nutrient assimilation, immune function, and neuro-behavioral processes.^14–16^ The disruption of a balanced GM composition, known as GM dysbiosis, directly impacts the host’s overall well-being, and has been linked to a spectrum of human diseases and conditions, including diabetes, asthma, obesity, Alzheimer’s and cancer.^17–21^

So far, *in vitro* and *in vivo* studies have predominantly concentrated on elucidating the mechanisms of oxidative stress ^22,23^ as a primary connection between human pesticide exposure and various chronic diseases. ^24,25^ Just recently, emerging research starts to shed light on the toxic effects of pesticides on the GM, while most of these studies are still mainly focused on observing qualitative or quantitative alterations in GM composition.^26–28^ Therefore, a systematic study of the metabolic responses of GM to various pesticides is critically needed, preferably with a selection of well-recognized, health-relevant, representative gut bacteria species and the adoption of universally accepted metabolic analysis approaches for GM. To address this critical need, we first introduced a microbiome-focused, integrated, mass-spectrometry-based metabolomics pipeline for the metabolic characterization of gut microbes in response to pesticide exposure. Then, by leveraging multi-omics investigations, our study discovered important interactions between representative gut bacteria species and well-recognized pesticide pollutants. Finally, we extended our investigation to a mouse model to verify the interactions between gut bacteria and the host following pesticide exposure and explore the consequential health implications of these interactions.

## Results

### Building a species-specific knowledge network of gut bacteria responses to pesticides

To assess the impacts of pesticides on human GM, we conducted a systematic analysis of interactions between eighteen compounds (comprising fifteen representative pesticides and three known pesticide metabolites) and seventeen representative human gut bacterial species (Table S1). This comprehensive interaction analysis yielded 306 bacteria-pesticide pairs, and four pesticides of particular concern were further examined for pesticide-dose-dependent growth inhibition or promotion (**Fig. 1a**). The selection of pesticides was based on their widespread agricultural use globally. Among the fifteen pesticides, eleven are still used across some areas of the world, despite some being identified as endocrine-disrupting chemicals (e.g., 4,4’-DDT, atrazine, and permethrin), persistent organic pollutants (e.g., chlorphyriphos, 4,4’-DDT), or prohibited or restricted in certain countries (**Fig. 1b**). Noteworthy variations in maximum residue limits (MRLs) for specific pesticides across different products and countries have raised concerns regarding potential health risks associated with these limits (**Extended Fig. 1a**). Meanwhile, to encompass a wide range of phylogenetic and metabolic diversity, we selected seven species from *Bacteroidetes*, seven species from *Firmicutes*, two species from *Actinobacteria*, and one species from *Proteobacteria* for initial growth inhibition/promotion experiments. As *Firmicutes* and *Bacteroidetes*, which collectively account for approximately 90% of the gut microbiota, are known to play significant roles in overall microbial diversity.^29^

**Fig. 1.**
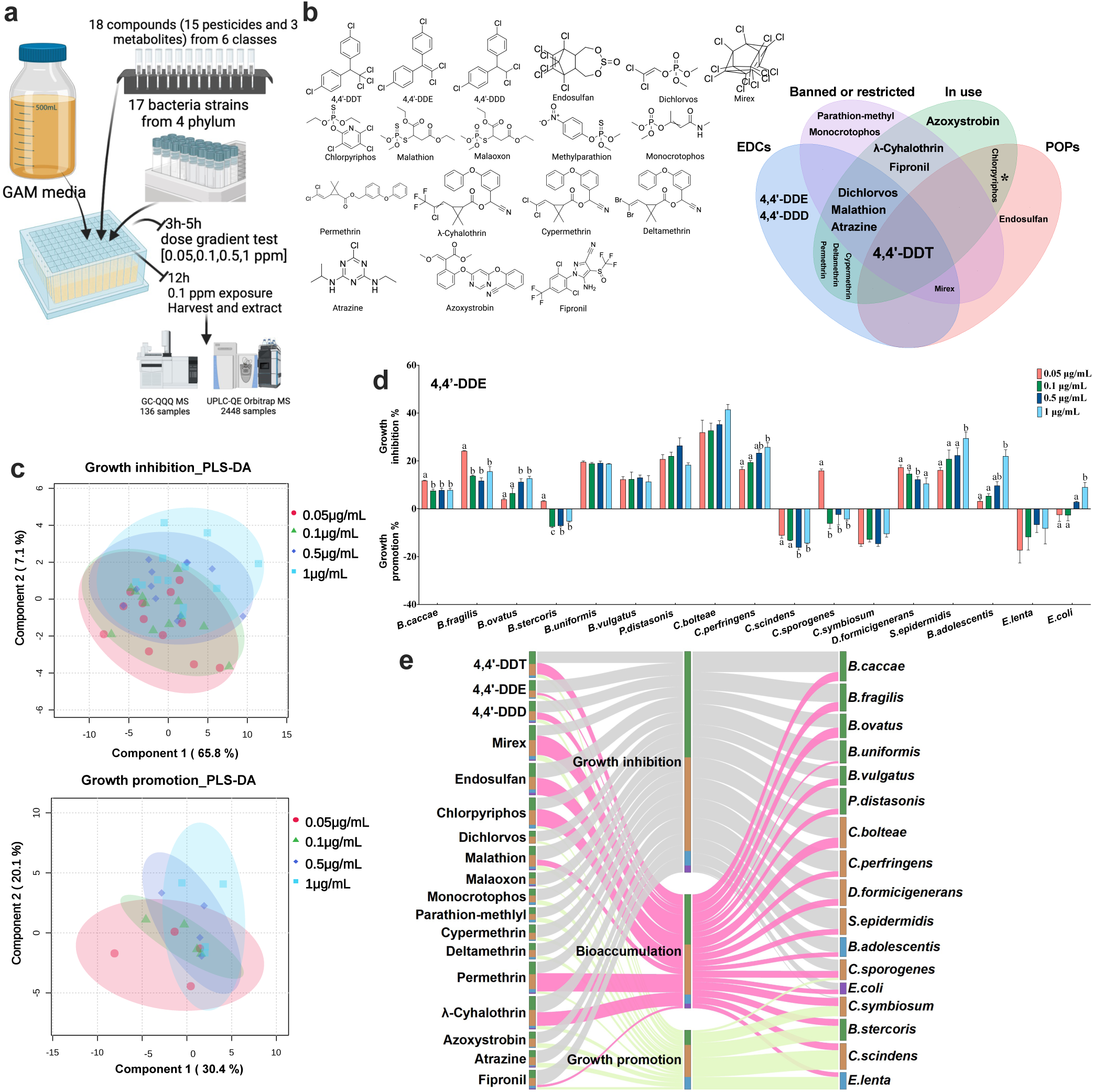
Gut bacteria accumulate pesticides without altering them. a, schematic of the assay. b, Structures of 18 compounds across 6 pesticide classes. c, pesticides elicit growth effects of gut microbiota at 0.05 μg/mL, 0.1 μg/mL, 0.5 μg/mL, and 1 μg/mL. d, 4,4’-DDE induced growth inhibition and promotion on 17 gut bacteria species under four concentrations. e, Bacteria-pesticide interaction network among growth inhibition/promotion and bioaccumulation in the study. Data are presented as mean ± SEM. p values were calculated by t-test, and p <0.05 represents statistically significant.

Based on the growth responses of these gut bacteria to pesticide exposure, they were classified into two clusters (Table S2): cluster I comprising pesticide-inhibited bacteria species, and cluster II, encompassing bacteria species that could be inhibited or promoted in a dose and pesticide-dependent manner (**Extended Fig. 1b**). The cluster I included *Bacteroides caccae*, *Bacteroides uniformis*, *Parabacteroides distasonis*, *Clostridium bolteae*, *Clostridium perfringenes*, *Dorea formicigenerans*, and *Staphylococcus epidermidis*. The cluster II included *Bacteroides fragilis*, *Bacteroides ovatus*, *Bacteroides vulgatus, Clostridium sporogenes*, *Bifidobacterium adolescentis*, and *Escherichia coli*, *Bacteroides stercoris*, *Clostridium scindens*, *Clostridium symbiosum*, and *Eggerthella lenta*. Furthermore, we conducted a partial least squares discriminant analysis (PLS-DA) to investigate the inhibition or promotion effects of 18 compounds on growth of 17 gut bacteria species (**Fig. 1c**). The PLS-DA plot effectively distinguished between different pesticide concentration groups, highlighting specific concentrations with different effects on bacterial growth. This analysis provided insights into the dose-dependent relationship, revealing the environmental impact of varying pesticide concentrations on gut bacterial growth. For instance, following exposure to 4,4’-DDE at four concentrations (**Fig. 1d**), the growth of 11 gut bacteria species was inhibited, 3 gut bacteria species were promoted, and 3 bacteria were shown conversely dose-dependent response to this pesticide. As the pesticide concentration increased, growth inhibition intensified in *B. ovatus*, *C. perfringens*, *S. epidermidis*, *B. adolescentis*, and *E. coli*, while growth promotion strengthened in *C. scindens*. Conversely, growth inhibition decreased in *B. caccae*, *B. fragilis*, *B. stercoris*, *C. sporogenes*, and *D. formicigenerans*. Within the concentration range of 0.05 μg/mL to 1 μg/mL, there were no changes observed at four concentrations for *B. uniformis*, *B. vulgatus*, *P. distasonis*, *C. bolteae*, and *E. lenta*. In summary, increasing pesticide concentration had varying effects on gut bacteria within a limited concentration range.

Furthermore, we investigated the capacity of gut bacteria to bioaccumulate pesticides after short-term (24-hour) exposure to 0.1 μg/mL of five organochlorine (4,4’-DDT, 4,4’-DDE, 4,4’- DDD, endosulfan, and mirex), two organophosphorous (chlorpyriphos and malathion) (Table S3), two pyrethroid (permethrin and λ-cyhalothrin) and one phenylpyrazole (fipronil) pesticides. The gut bacteria-pesticide interaction network (**Fig. 1e**) indicated that the growth of most gut bacteria was inhibited by all pesticides, while only a few was promoted. However, our results indicated that all gut bacteria species can bioaccumulate selected pesticides, and the ability of bioaccumulation is pesticide type-dependent. These results quantitatively confirmed the long-standing speculations that pesticide induced growth inhibition or promotion on gut bacteria. Our data also supported that pesticides disrupt GM composition, while bacteria accumulation of pesticide increases the risk of prolonged pesticide residue in the host.

### Mapping the pesticide-GM-metabolites (PMM) interactions

It is often assumed that gut bacteria responses to pesticide are associated with the changes of gut bacterial metabolites, thereby regulating host health. However, experimental evaluation of these responses metabolically has not been done, to fill the gap of knowledge, we conducted high-throughput metabolomics analyses to elucidate changes in endogenous metabolites in gut bacteria species after pesticide exposure *in vitro*. To elucidate the mechanisms underlying the changes of microbial metabolism in response to pesticides, we first profiled over 472 (**Table S4**) metabolites from the pesticides-bacteria network interaction experiments, and summarized these metabolic responses of pesticide-inhibited or promoted bacteria (**Fig. 2a** and **Table S5**). We found six highly pesticide-sensitive gut bacteria exhibiting the most significant changes in terms of the number of metabolites impacted. These species include *B.ovatus*, *B.uniformis*, *D.formicigenerans*, *B.stercoris*, *C.symbiosum,* and *B.adolescentis* (**Fig. 2a**). We further mapped the significant metabolic alterations into 40 metabolic pathways (**Fig. 2b** and **Table S6**), which encompassing amino acid metabolism (**Extended Fig. 3**), carbohydrate metabolism, cofactors and vitamins metabolism, nucleotide metabolism, and sulfate/sulfite metabolism (**Extended Fig. 4**). Among these pathways, the top 5 of most commonly affected pathways included pyrimidine metabolism, purine metabolism, arginine and proline metabolism, lysine degradation, and phenylalanine and tyrosine metabolism. It is expected that the gut bacterial metabolites from those affected pathways could further serve as molecule signals influencing host health, as reported in other studies ^30,31^. Notably, we observed that pesticides directly impact tryptophan metabolism (**Fig. 3a**), propanate metabolism (**Fig. 3b**) and butyrate metabolism (**Fig. 3c**) in several gut bacteria species, therefore leading to the dysregulated production of indole and its derivatives (indoles), and SCFAs (**Extended Fig. 3-4**) which will in turn modulate human metabolism and impact human health.^32^ It is also interesting to note that pesticides can affect the same pathways in different gut bacteria species by inducing the dysregulated production of distinct metabolites within those pathways (**Fig. 3a-c** and **Table S3-4**). For instance, pesticides influenced tryptophan metabolism in ten gut bacteria species, and each of these impacted bacteria exhibits distinct pattern of pesticide-sensitive metabolites (**Fig. 3a** and **Extended Fig. 3**); In *B.ovatus*, decreased L-tryptophan, N1-acetyl-N2- formyl-5-methoxykynuramine, 6-hydeoxymelatonin, anthranilate, trans-3-indoleacrylic acid, and increased indole, indole-3-acetic acid, indole-3-acetly-L-alanin and 2-aminomuconic acid were observed when exposed to the tested pestcides, while we found only increased 2-amino-3,7- dideoxy-D-threo-hept-6-ulosonate in *B.vulgatus* with the same exposure experiments. Meanwhile, all pesticides can affect the tryptophan metabolism in *B.ovatus*, but only dichlorvos can affect it in *B.vulgatus* (**Fig. 3a** and **Extended Fig. 3**). Through Pearson correlation analysis of pesticide induced bacterial growth perturbation, and the significant changes in the number of metabolites, we identified several unique metabolites that are sensitive to the pesticide exposure in a bacteria-specific manner (**Fig. 3d**). In the study, we identified pesticide-sensitive gut bacteria species and specific pesticide-gut bacteria pairs that regulate important microbial metabolic pathways. This complexity underscores the challenges in identifying biomarkers in gut bacteria exposed to pesticides and highlights the importance of metabolomics analysis in understanding these effects. Through our high-throughput data integration, we have delineated important network information on PMM interactions, and provided a rich dataset for research into the pathogenic mechanisms of pesticides’ impact on human health.

**Fig. 2.**
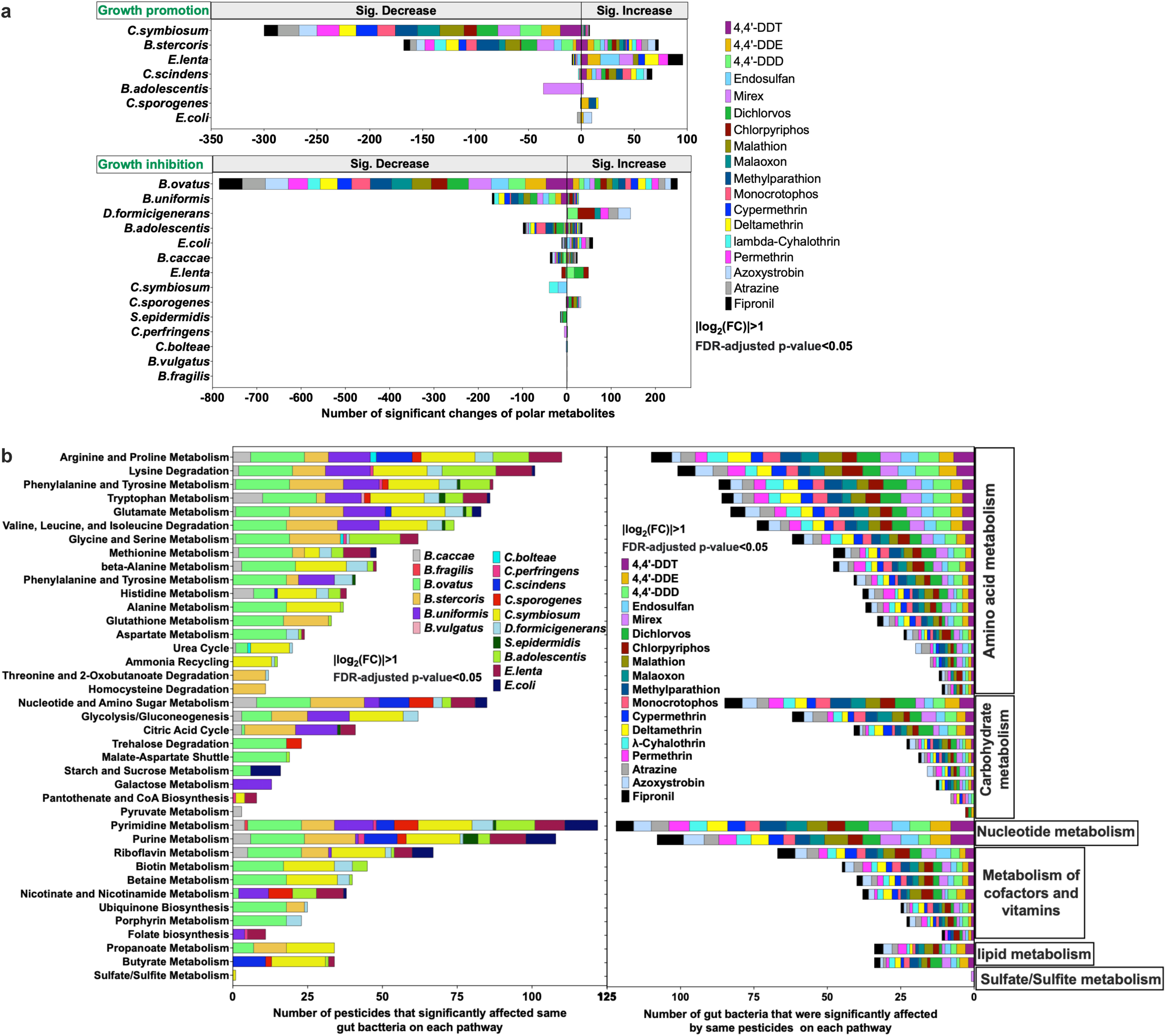
pesticide exposure led to broad and systemic changes in metabolites across gut microbiota. a, number of significant changes in metabolites for bacteria species with growth promotion or inhibition after exposure to 18 compounds. b, number of pesticides that significantly affected polar metabolic pathways for each bacteria strain. Data are presented as |log2(FC)|>1 and FDR-adjusted p-value <0.05 (a-d). FC, fold change; FDR, false discovery rate.

**Fig. 3.**
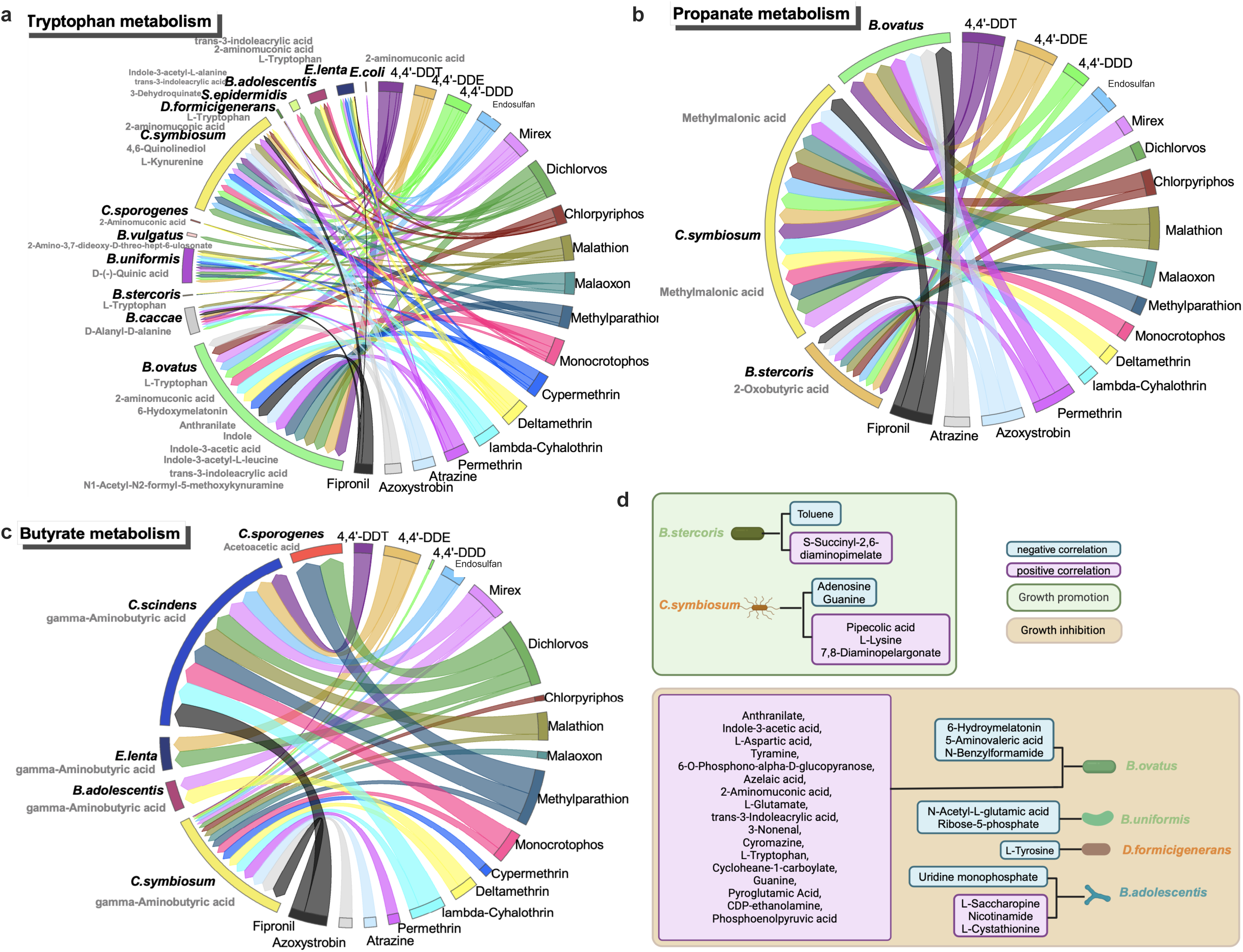
Network interaction that pesticide exposure led to changes in metabolites across gut microbiota. a, network interaction that pesticides affected tryptophan metabolism for ten bacteria species by different metabolites. b, network interaction that pesticides affected propanate metabolism for three bacteria species by different metabolites. c. network interaction that pesticides affected butyrate metabolism for five bacteria species by different metabolites. d, Pearson correlation analysis between growth promotion/inhibition and significant changes of metabolites. Data are presented as |log2(FC)|>1 and FDR-adjusted p-value <0.05 (a-d). FC, fold change; FDR, false discovery rate.

### Discovering lipid molecule changes in PMM at multiple levels

GM is increasingly recognized as an endocrine organ, producing secretions that can influence the body through the bloodstream or lymphatic system. ^33^ And the GM has the capacity to transform and synthesize bioactive lipids with structural and signaling functions, impacting host metabolism and immunology.^34^ While our study included several known endocrine disrupting pesticides, it is plausible that undefined endocrine disruptors could also modulate the production of bioactive lipids by the GM. Therefore, we further conducted a comprehensive lipidomics analysis to investigate how pesticides affect GM lipid molecule changes, and reported the pesticides-GM lipids interaction network here from various levels such as lipid categories, classes, chain lengths of fatty acyl, and saturation status (**Fig. 4a** and **Table S7-8**). Our findings revealed that pesticides induced most significant changes in the quantity of lipid molecules in pesticide-promoted bacteria compared to pesticide-inhibited bacterial species (**Fig. 4b** and **Extended Fig. 6a**); Specifically, pesticides significantly decreased the detected level of many lipids in *B.stercoris* and significantly increased the detected level of hundreds of lipids in *C.symbiosum* and *C.scindens*, while causing much fewer lipid changes in pesticide-inhibited *B.ovatus* (**Fig. 4b**). Furthermore, we found that the most significant changes in lipid class quantities were concentrated in several bacterial species, including *B.stercoris*, *C.scindens*, and *C.symbiosum*. The results suggested that the lipid metabolism of these gut bacteria are highly sensitive to pesticide exposure. However, no single pesticide was identified as unique cause of these changes; rather, all pesticides consistently influenced lipid class alterations across each bacterial species (**Fig. 4c**). Overall, the most significant changes of lipids in gut bacteria occurred in glycerophospholipids (GPs) category (**Fig. 4c**), which are expected as they are known to be abundant in bacterial cell membranes and therefore are generally at the frontier of attack by pesticides.^34^ Diving into the lipid classes level, we further observed that pesticides significantly influenced acylcarnitine (CAR, CAR 20:0) from FAs, diacylglycerol (DG, such as DG 16:1_17:1) and ether-linked diacylglycerol (EtherDG, such as DG O-16:2_17:1) from GLs, ceramide non-hydroxyfatty acid-dihydrosphingosine (Cer_NDS, such as Cer 17:0; 2O/15:0) from SPs, and ether-linked lysophosphatidylglycerol (EtherLPG, such as PG O-17:1_16:0) from GPs in many gut bacteria and in particular in *B.stercoris*, *C.symbiosum*, and *C.scindens* (**Fig. 4c** and **Extended Fig. 5a**).

**Fig. 4.**
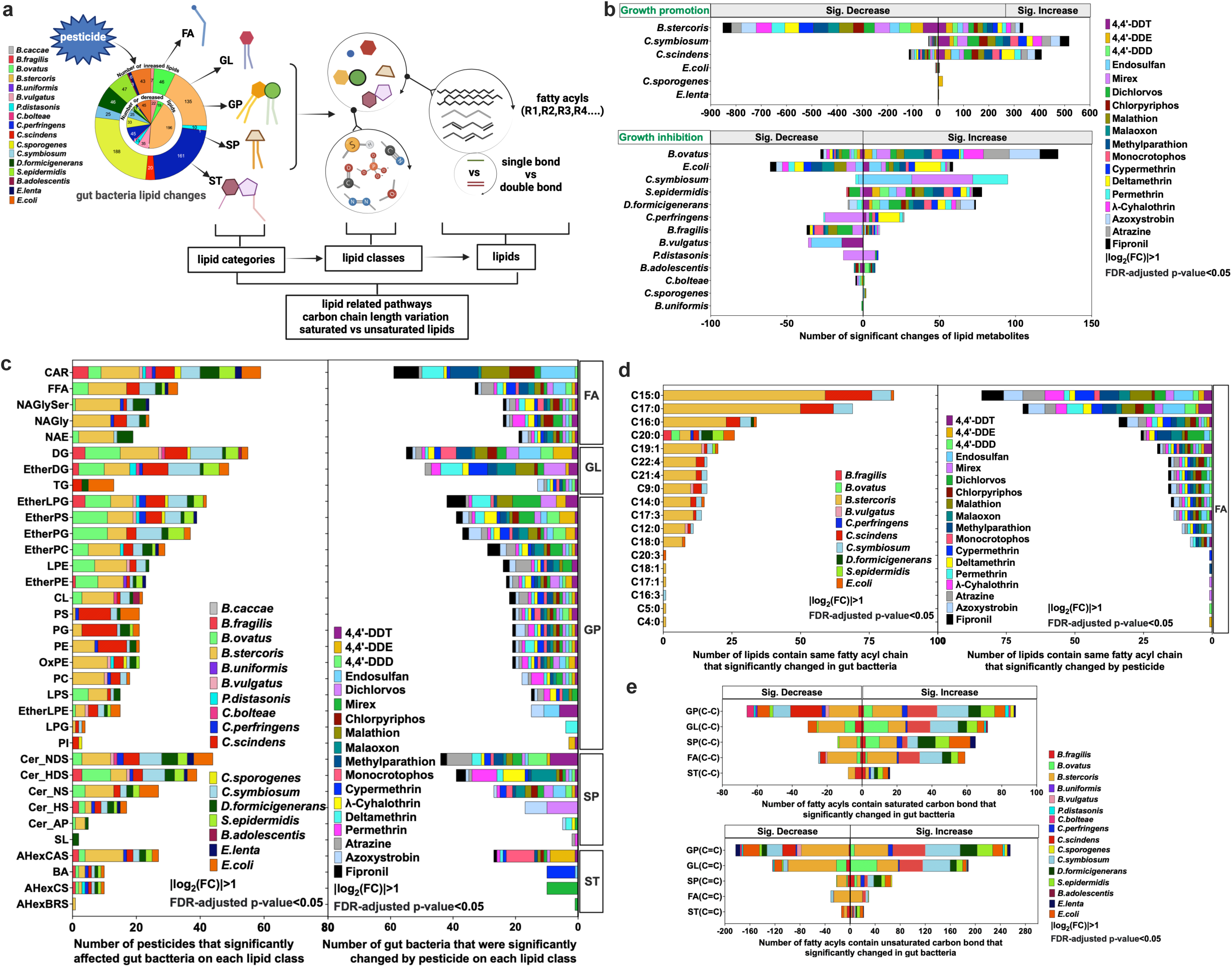
pesticide exposure induces extensive and systemic changes in lipids across gut bacteria species. workflow for lipid metabolites analysis including lipid categories, lipid classes and lipids. a. number of significant changes in lipid metabolites for bacteria species with growth promotion or inhibition after exposure to 18 compounds. b. number of pesticides or gut bacteria species that were significantly changed on each lipid classes. c. Number of lipids contain same fatty acyl chain that significantly changed for FA category. d. Number of lipid carbon bond that significantly changed in gut bacteria by pesticide exposure. Data are presented as |log2(FC)|>1 and FDR-adjusted p-value <0.05 (a-d). AHexBRS, Acylhexosyl brassicasterol; AHexCAS, Acylhexosyl campesterol; AHexCS, Acylhexosyl cholesterol; BA, bile acid; CAR, Acylcarnitine; Cer, ceramide; Cer_AP, Ceramide alpha-hydroxy fatty acid-phytospingosine; Cer_HS, Ceramide hydroxy fatty acid-sphingosine; Cer_HDS, Ceramide hydroxy fatty acid-dihydrosphingosine; Cer_NDS, Ceramide non-hydroxyfatty acid-dihydrosphingosine; Cer_NS, Ceramide non-hydroxyfatty acid-sphingosine; CL, cardiplipin; DG, diacylglycerol; EtherDG, Ether-linked diacylglycerol; EtherLPE, Ether-linked lysophosphatidylethanolamine; EtherLPG, Ether-linked lysophosphatidylglycerol; EtherPC, Ether-linked phosphatidylcholine; EtherPE Ether-linked phosphatidylethanolamine; EtherPG, Ether-linked phosphatidylglycerol; EtherPS, Ether-linked phosphatidylserine; FA, fatty acyl; FFA, fatty acid; FC, fold change; FDR, false discovery rate; HexCer, hexosylceramide alpha-hydroxy fatty acid-dihydrosphingosine; MGDG, Monogalactosyldiacylglycerol; GL, Glycerolipid; GP, Glycerophospholipid; LPE, ether-linked lyspphosphatidylethanolamine; LPG, Lysophosphatidylglycerol; LPS, Lysophosphatidylserine; NAE, N-acyl ethanolamines; NAGly, N-acyl glycine; NAGlySer, N-acyl glycyl serine; NAOrn, N-acyl ornithine; NATau, N-acyl taurine; oxFA, Oxidized fatty acid; oxTG, Oxidized triglyceride; oxPS, Oxidized phosphatidylserine; PC, phosphatidylcholine; PE, phosphatidylethanolamine; PEtOH, Phosphatidylethanol; PI, Phosphatidylinositol; PG, phosphatidylglycerol; PS, phosphatidylserine; SL, sulfonolipid; SM, Sphingomyelin; SP, Sphingolipid; SSulfate, Sterol sulfate; ST, sterol lipid; TG, triacylglycerol.

We also discovered that, in FAs, the most significant changes in the quantity of fatty acyl chains occurred in *B.stercoris*, while saturated C20:0 significantly changed in 9 bacteria species (**Fig. 4d**). Therefore, the saturated fatty acid C20:0, arachidic acid, may be further considered as a sensitive bioindicator for gut bacteria following pesticide exposure. Compared to even numbered-lipids in mammals, bacterial lipids embrace greater diversity with both odd- or even-numbered fatty acyl chains reported in previous studies,^35^ such as C15/ C17/ C19 vs C16/ C20/ C22 in the FAs (**Fig. 4d**), glycerols (GLs) (**Extended Fig. 6b**), GPs (**Extended Fig. 6c**), and SPs (**Extended Fig. 6d**). After pesticides exposure, significant changes in the quantity of fatty acyl chains were observed in C15:0, C16:0, C17:0, and C17:1 in FAs, GLs, GPs, SPs (**Fig. 4d** and **Extended Fig. 6b-d**), while C28:1 and C9:0 were affected in STs (**Extended Fig. 6e**); Odd chain fatty acids (OCFAs), C15:0, C17:0 and C17:1 can be enlongated to very-long-chain FAs (VLCFAs) or can be derived from these VLCFAs. ^36^ Their chain can be shortened, yielding propionyl-coenzyme A (CoA) to replenish the citric acid cycle, and the concentrations of C15:0 and C17:0 were associated to multiple disease risk, and involved in discussions of biomarker identification or treatment pathway. ^37^ Furthermore, C 9:0 can alter small intestine neuroendocrine tumor phenotype ^38^ and regulate epithelial immunological barrier function. ^39^ Meanwhile, those changes for each lipid category were identified to be associated with specific gut bacteria. For example, pesticides induce most changes of fatty acyls chains in *B.stercoris* across all lipid categories. However, more changes in *C.symbiosum* and *C.scindens* across GLs and GPs, in *B.stercoris*, *D.formicigenerans*, *S.epidermidis*, and *E.coli* across SPs and STs, and *B.fragilis*, *B.stercoris*, *B.ovatus*, and *E.coli* in STs were observed. Furthermore, it is well-known that lipid A (endotoxin), a component of lipopolysaccharide (LPS), is a glucosamine-based phospholipid that typically contains C14, C16 and C18 hydroxy acyl chains in most Gram-negative bacteria. ^40–42^ While significant changes in C14:0, C16:0 and C18:1 in GPs and GLs in gut bacteria, especially for Gram-negative bacteria *B.stercoris* after pesticide exposure (**Extended Fig. 6b-c**) were also observed in our study. We speculated that pesticides can disrupt the levels of those lipids in certain gut bacteria, such as *B.stercoris*, leading to a potential dysregulation of lipid A or LPS, ^43^ and indirectly affecting the host immune system. ^44^ Additionally, pesticides predominantly affected lipids with carbon bond across all lipid categories (**Fig. 4e**, and **Extended Fig. 6f**) in *B.stercoris*, *C.symbiosum*, *C.scindens*, and *E.coli*. The substantial increase in the number of saturated C-C bonds suggests that gut bacteria may be adapting to pesticide exposure through an oxidative stress response.^45,46^

### Expanding PMM study in a mouse model: discovering the *B.ovatus* vs 4,4’-DDE interactions *in vivo*

Considering the extensive effects of pesticides on gut bacteria and metabolite levels *in vitro*, we further selected a representative pesticide-gut bacteria pair to explore the pesticide-gut bacteria-host interactions *in vivo*. Among many of the pesticide-inhibited bacteria, *B.ovatus* (a Gram-negative bacterium) was selected for the *in vivo* study based on is sensitive metabolic responses to the pesticide perturbation, as evidenced by the largest number of significantly changed metabolites detected (**Extended Fig. 7a**). Meanwhile, organochlorine pesticides are known to interfere with inflammation responses in the host,^47,48^ and 4,4’-DDE was chosen due to its profound impact on *B.ovatus* metabolism (**Fig. 5a,b** and **Extended Fig. 7b**). The *in vivo* study comprised three mouse groups: named as ABX (only antibiotic-treated), Control (antibiotic-treated and 4,4’-DDE exposure), and BO (antibiotic-treated, *B.ovatus* transplantation and 4,4’-DDE exposure) groups. After 4,4’-DDE exposure in the Control and BO group, notable changes in microbial composition were observed (**Fig. 5c**) with varying levels of 4,4’-DDE detected in mouse organs and tissues (**Fig. 5d**).

**Fig. 5.**
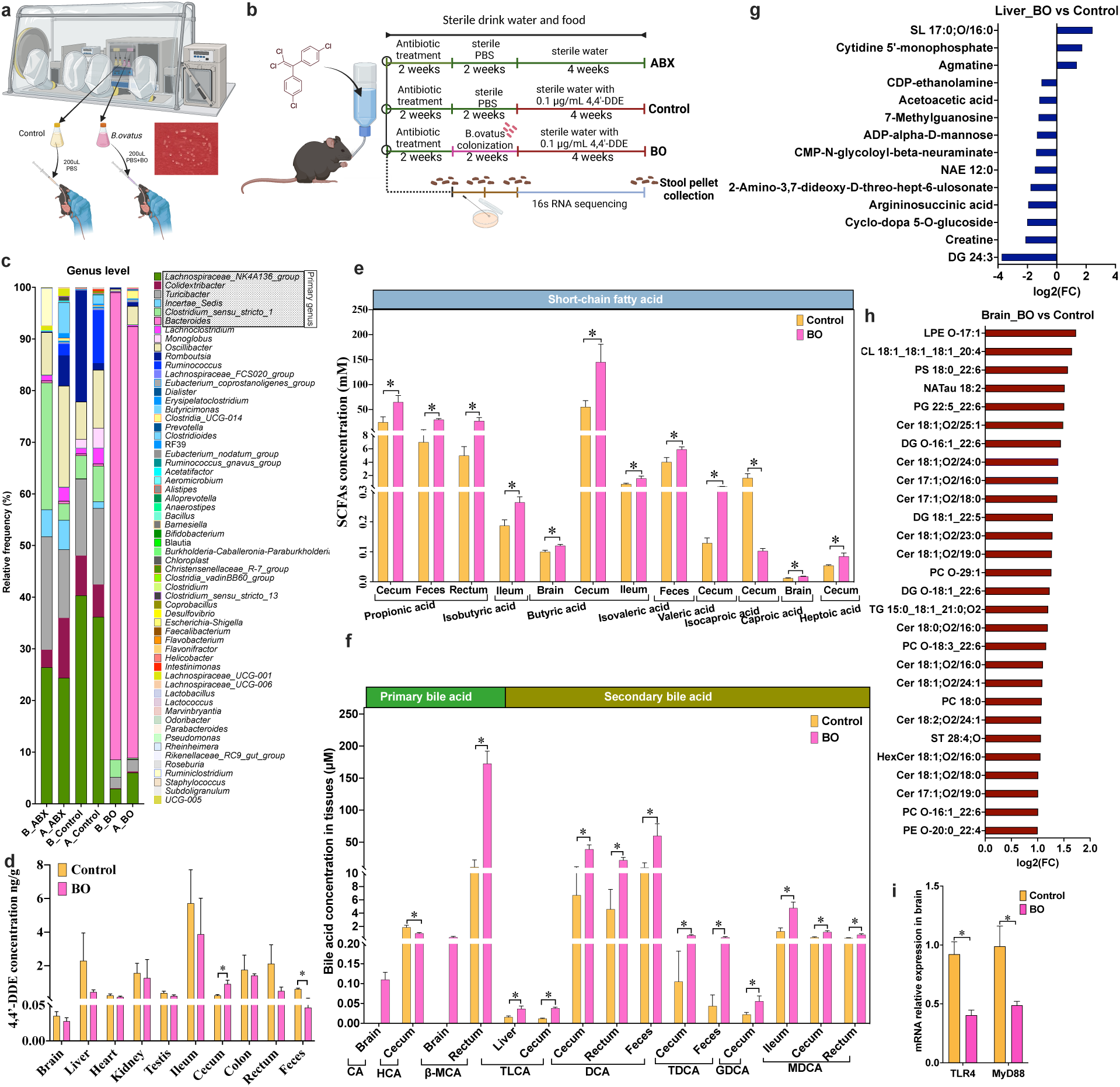
metabolic changes induced by 4,4’-DDE in *B.ovatus* transplanted C57BL/6 mice. a, cultivation of *B. ovatus* in anaerobic chamber and transplantation into C57BL/6 mice via stomach gavage. b, schematic representation of *B. ovatus* transplantation and 4,4’-DDE exposure in three groups of C57BL/6 mice. c, changes in gut microbiota composition before and after 4,4’-DDE exposure over four weeks in three groups. d, detection of 4,4’-DDE in organs and tissues of mice in the Control and BO groups. e, significant changes in SCFAs in organs and tissues of mice using a targeted method at the end of experiment in the Control and BO groups. f, significant changes of BAs in organs and tissues of mice using targeted method at the end of experiment in the Control and BO groups. g, untargeted metabolomics and lipidomics revealing significant metabolite changes (|log2(FC)|>1 and p-value <0.05) in the liver of the BO group compared to the Control group at the end of experiment. h, untargeted metabolomics and lipidomics revealing significant metabolite changes (|log2(FC)|>1 and p-value <0.05) in the brain of the BO group compared to the Control group at the end of experiment. i, significant changes in mRNA relative expression of receptors from inflammation signaling pathway in the brain at the end of experiment in the Control and BO groups. Data (d-f, i) are presented as mean ± SEM. p values were calculated by t-test, and p <0.05 represents statistically significant. SCFAs, short-chain fatty acids; BAs, bile acids; CA, cholic acid; HCA, hyocholic acid; β-MCA, β-muricholic acid; TLCA, taurolithocholic acid; DCA, deoxycholic acid; TDCA, taurodeoxycholic acid; GDCA, glycodeoxycholic acid; MDCA, murideoxycholic acid; FC, fold change; SL, sulfonolipid; CDP, cytidine-5’-diphosphate; ADP, adenosine-5’- diphosphate; CMP, cytidine monophosphate; NAE, N-acyl ethanolamines; DG, diacylglycerol; LPE, ether-linked lyspphosphatidylethanolamine; CL, cardiplipin; PS, phosphatidylserine; NATau, N-acyl taurine; PG, phosphatidylglycerol; Cer, ceramide; PC, phosphatidylcholine; TG, triacylglycerol; ST, sterol lipid; HexCer, hexosylceramide alpha-hydroxy fatty acid-dihydrosphingosine; PE, phosphatidylethanolamine; TLR4, musculus toll-like receptor 4; MyD88, myeloid differentiation factor 88.

Furthermore, our results indicated that during the 4,4’-DDE exposure, the addition of *B.ovatus* in the BO group significantly increased SCFAs and secondary BAs in the liver, brain and intestine of mice compared to the control group, which may potentiate anti-inflammatory effects in these organs ^49–51^ (**Fig. 5e**-**f** and **Extended Fig. 7c-d**). Meanwhile, the addition of *B.ovatus* in the BO group also decreased branched-SCFA isocaproic acid and primary bile acid hyocholic acid (HCA) in the cecum. Considering SCFAs and BAs could act via the gastrointestinal tract to modulate the host immune system, ^52–54^ subsequently, we performed analyses that focused on metabolism of the liver and brain, as well as the Toll-like receptor 4 (TLR4)/nuclear factor κB (NF-κB) inflammatory signaling pathway. In the liver, significant changes in amino acid metabolism were detected (**Fig. 5g** and **Extended Fig. 7e-f**); conversely, in the brain, 28 significantly increased metabolites were all lipids from GLs, GPs, SPs, and STs (**Fig. 5h** and **Extended Fig. 7g-h**) in the BO group. Additionally, significantly lower levels of TLR4 and myeloid differentiation factor 88 (MyD88) to attenuate inflammation were observed in the BO group compared to the Control group (**Fig. 5i** and **Extended Fig. 7i**). Here, most SCFAs and BAs from *B.ovatus*, as well as lipids (GLs, GPs, SPs, and STs) from brain, were negatively correlated with the receptors from the TLR4/ NF-κB pathway (**Extended Fig. 7j**). Recall from the earlier results that the 4,4’-DDE also significantly increased the level of lipids (FAs, GLs and GPs) in *B.ovatus in vitro* (**Extended Fig. 7k**), these findings collectively suggest that pesticides can affect the lipid metabolism of both gut bacteria and the host to regulate the level of lipid molecules in host and attenuate inflammation.

## Discussion

Previous studies have suggested that GM plays a crucial role in pesticide breakdown and intricately mediate the impacts of pesticides on human health. ^55,56^ This relationship is partially characterized by the alterations in GM composition ^57–60^ as well as changes in host metabolites ^61,62^ following pesticide exposure. Recognizing that the GM functions and metabolism may also be inadvertently impacted by pesticide, we aimed to assess the toxicity induced by pesticides on representative gut bacteria and its subsequent effects on host health via comprehensive metabolomics analyses. In our study, we observed that pesticides have the ability to perturbate the growth of gut bacteria in a bacteria-pesticide specific manner even at low concentration range from 0.05 μg/mL to 1 μg/mL (**Fig. 6a**). This finding provides compelling evidence for the previously observed imbalance in gut bacteria communities following pesticide exposure. ^63–65^ Furthermore, the existence of bacteria capable of accumulating pesticides under pesticide exposure directly contributes to long-term pesticide residue or endocrine disruption in the host. These findings highlight the intricate relationship between pesticides and the GM, emphasizing the need for further research to understand the implications of pesticide exposure on human health.

**Fig. 6.**
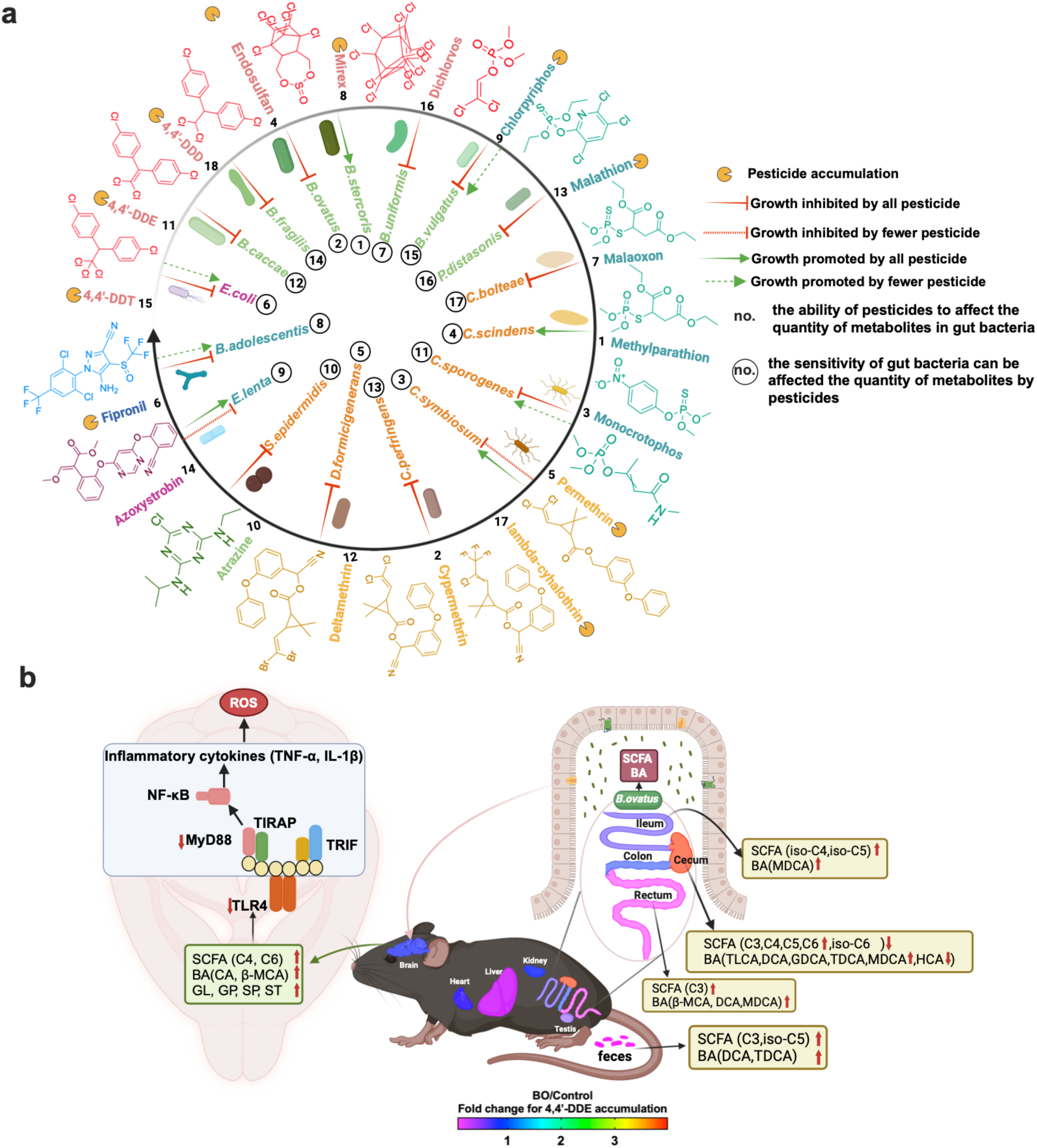
comprehensive network interaction analysis between gut microbiota and pesticide using *in vitro* and *in vivo* models. a, pesticide impact on gut microbiota: growth inhibition/promotion and bioaccumulation. b, metabolic changes induced by 4,4’-DDE in C57BL/6 mice: targeted and untargeted analysis. SCFA, short-chain fatty acid; C3, propionic acid; C4, butyric acid; iso-C4, isobutyric acid; C5, valeric acid; iso-C5, isovaleric acid; C6, caproic acid; BA, bile acid; CA, cholic acid; HCA, hyocholic acid; β-MCA, β-muricholic acid; TLCA, taurolithocholic acid; DCA, deoxycholic acid; TDCA, taurodeoxycholic acid; GDCA, glycodeoxycholic acid; MDCA, murideoxycholic acid; GL, glycerolipid; GP, glycerophospholipid; SP, sphingolipids; ST, sterol lipids; IL-6, interleukin 6; IL-1β,interleukin 1 β; MyD88, myeloid differentiation factor 88; NF-κB, nuclear factor kappa-light-chain-enhancer of activated B cells; TLR4, musculus toll-like receptor 4; TNF-α, tumor necrosis factor alpha; TRIF, TIR-domain-containing adapter-inducing interferon-β.

Concomitant with changes in growth status, the metabolic susceptibilities of tested gut bacteria were measured and evaluated in this pesticide-gut bacteria interaction network study. Gut bacteria are known to influence the levels of SCFAs ^66^, BAs ^67^, trimethylamine (TMA) ^68^, serotonin (5-HT) ^69^, and LPS ^70^, all of which are critical for host health. Moreover, we identified several pesticide-sensitive gut bacteria, suggesting that a diverse group of pesticides can uniquely target susceptible metabolites in different microbial metabolic pathways. Additionally, biochemical characterizations of lipid categories, classes, fatty acyl chain lengths, and lipid saturation levels in gut bacteria post pesticides exposure were performed in our study, and we observed many bacterial lipids affected by pesticide exposure. And changes in fatty acyls in the pesticide-sensitive gut bacteria following pesticide exposure can be used for monitoring pathogenesis. While the composition and function of the GM in the gut could be affected, variations among GM in human populations and their uncertain functions for human health present challenges in establishing microbial reference communities for pesticide toxicity evaluations. Therefore, our study not only provided a robust data resource but also standardized experimental approaches to identify the growth perturbations caused by multiple pesticides on various representative human gut bacteria. Pesticide exposure not only affects bacterial growth conditions but also influences their metabolism, potentially disrupting the crucial interaction between the microbiota and the host. While not all GM perturbations lead to adverse effects, our study aimed to conduct a risk assessment of GM-mediated pesticide exposure on the host. Our analysis of *B. ovatus* demonstrated its ability to enhance the pesticide elimination in many major organs or tissues in the host (**Fig. 6b**). Concurrently, transplanted *B. ovatus* increased the levels of SCFAs and BAs in brain, which is in alignment with other observations ^71,72^. The elevation of lipids in the brain, including SCFAs, BAs, GLs, GPs, SPs, and STs, in turn regulates inflammation-related signaling pathways, as demonstrated in our detected downregulation of mRNA expression of TLR4 and MyD88. These connections suggested a *B.ovatus*-mediated gut-brain axis after 4,4’-DDE exposure. In summary, our current dataset, which includes strain-specific pesticide-metabolic profiles, offers a valuable resource for the comparative identification of biomarkers and the development of preventive strategies in understanding the interplay between gut bacteria and the host following pesticide exposure. Moreover, these data, along with the associated methodology, can serve as a direct reference or a readily deployable platform for enhancing the detection of microbiome-dependent metabolite perturbation in biological samples after pesticide exposure.

### Limitations of the study

Our study aimed to enhance the fundamental understanding of pesticide exposure on gut bacteria, both *in vitro* and *in vivo*, in a generally healthy population. Given the complexity and diversity of the human GM, alongside the variability in environmental contaminants, we focused on a subset of microbiota and pesticides. This approach leaves many potential interactions unexamined. Currently, our research has concentrated solely on metabolomics and lipidomics outcomes. To gain a more comprehensive understanding, future studies should incorporate additional omics approaches to screen for functional genes and their related metabolites. The next phase of our research will involve conducting animal model experiments on specific diseases, targeting pesticide-bacteria pairs that produce particular metabolites. For example, after pesticide exposure, the lipid and glucose metabolism of GM were significantly affected; we will further investigate the health impacts on the host through disease mouse model (such as obesity model) to further understand the contribution of the interactions between pesticides and microbiota to the disease pathogenesis. This will finally contribute to the regulation of pesticide residues based on human health and the prevention of diseases.

## STAR METHODS

Detailed methods are provided in the online version of this paper and include the following:

- **KEY RESOURCE TABLE**
- **RESOURCE AVAILABILITY**

Lead contact
Materials availability
Data and code availability
- **METHOD DETAILS**

Data reporting
Growth conditions
Pesticide dose-dependent growth inhibition
Pesticide bioaccumulation detection and metabolism evaluation
Mouse experiment
Mouse samples preparation for pesticide analysis
Mouse samples preparation for SCFAs, BAs, and metabolomics analysis
Mouse samples preparation for lipidomics analysis
GC- QQQ MS methods
UPLC-QE Orbitrap MS methods Transcriptomics analysis
16s RNA sequencing
Data analysis

Masshunter quantitative analysis Compound discover analysis MS-DIAL analysis
Statistics and reproducibility Data availability
Code availability

## SUPPLEMENTAL INFORMATION

Supplemental information can be found online.

## ACKNOWLEDGEMENTS

This study was supported by the National Institute of General Medical Sciences of the National Institutes of Health (R35GM133510).

## AUTHOR CONTRIBUTIONS

Conceptualization, L.C. and J.Z.; Methdology, L.C., J.Z., H.Y. S.D., and Y.W.; Investigation, L.C., C.G, H.Z, A.G., S.Z., D.W., M.H., C.J., and X.W.; Writing-Orignial Draft, L.C. and J.Z.; Visualization, L.C.; Funding Acquisition, J.Z..

---

**Extended Fig.1.**
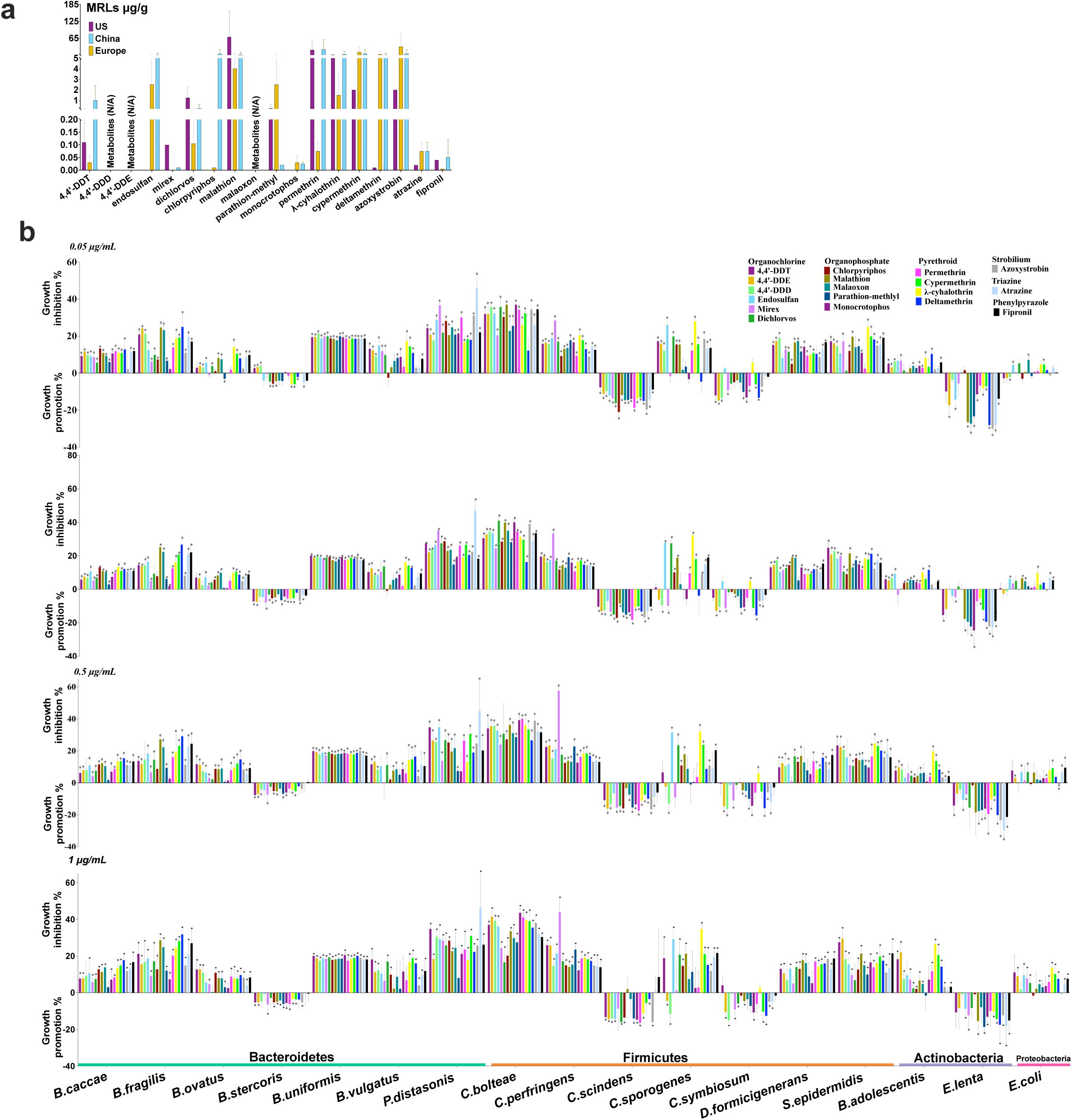
Selection of 18 compounds and their impact on gut microbiota growth inhibition/promotion. a, pesticides usage status and potential toxic effects on organisms. b, pesticides and their metabolites induced growth inhibition/promotion for 17 gut bacteria strains across four concentrations (0.05 μg/mL, 0.1 μg/mL, 0.5 μg/mL, and 1 μg/mL). Data (d) are presented as mean ± SEM (n=4). p values were calculated by t-test, and p <0.05 represents statistically significant.

**Extended Fig.2.**
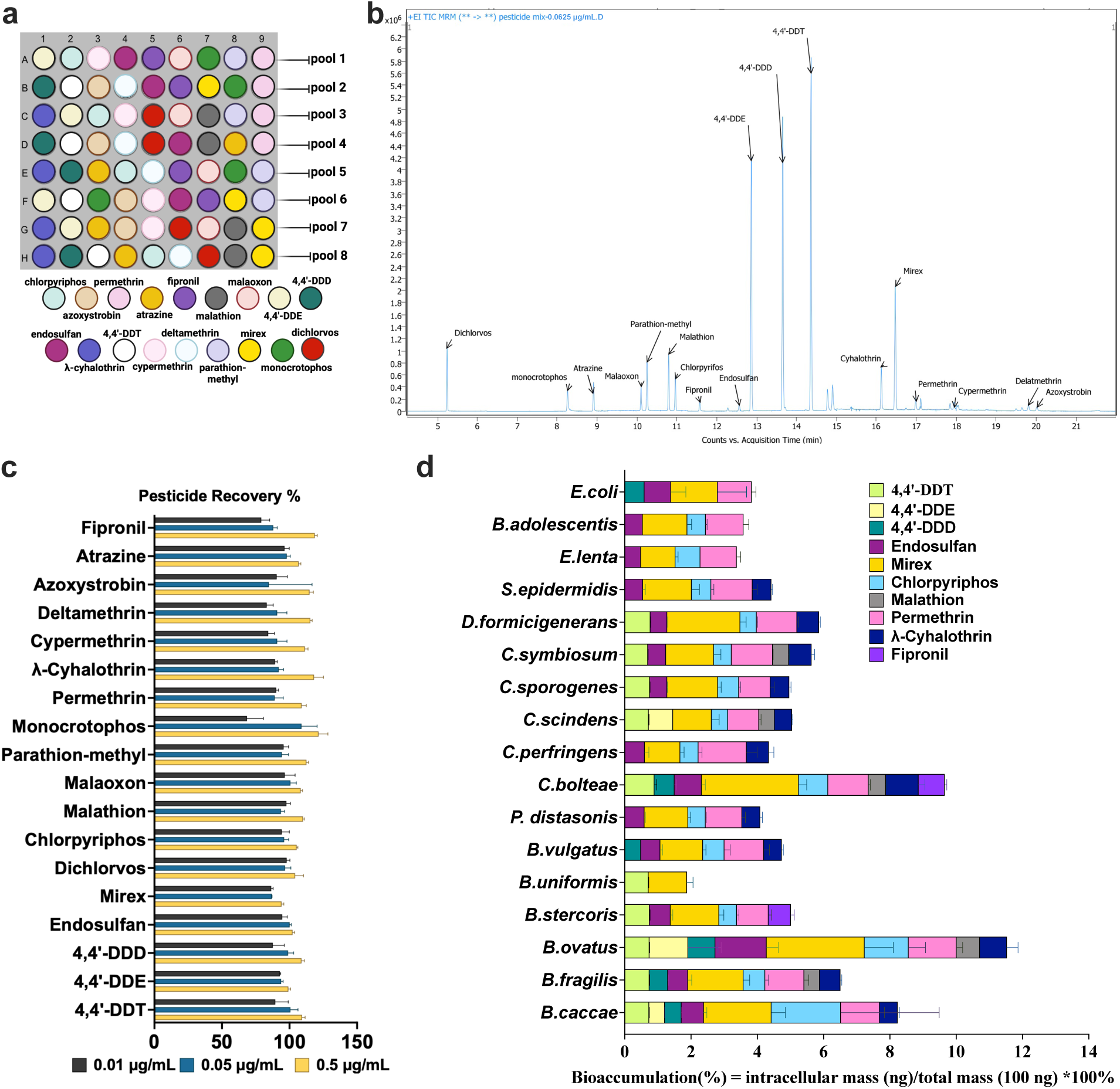
Detection of 18 compounds in gut microbiota by GC-QQQ MS. a, a combinatorial pooling strategy to allocate 18 compounds into 8 pools to detect bioaccumulation. b, chromatogram of standard mix of 18 compounds with retention time by GC-QQQ MS. c, pesticide recoveries in GAM media under three concentrations (10 ng/mL, 50 ng/mL, and 500 ng/mL) d, bioaccumulation of 10 out of 18 compounds by gut bacteria strains in 24 hours. Data (d) are presented as mean ± SEM (n=4). p values were calculated by t-test, and p <0.05 represents statistically significant. GC-QQQ MS, gas chromatograph coupled triple quadruple mass spectrometry.

**Extended Fig.3:**
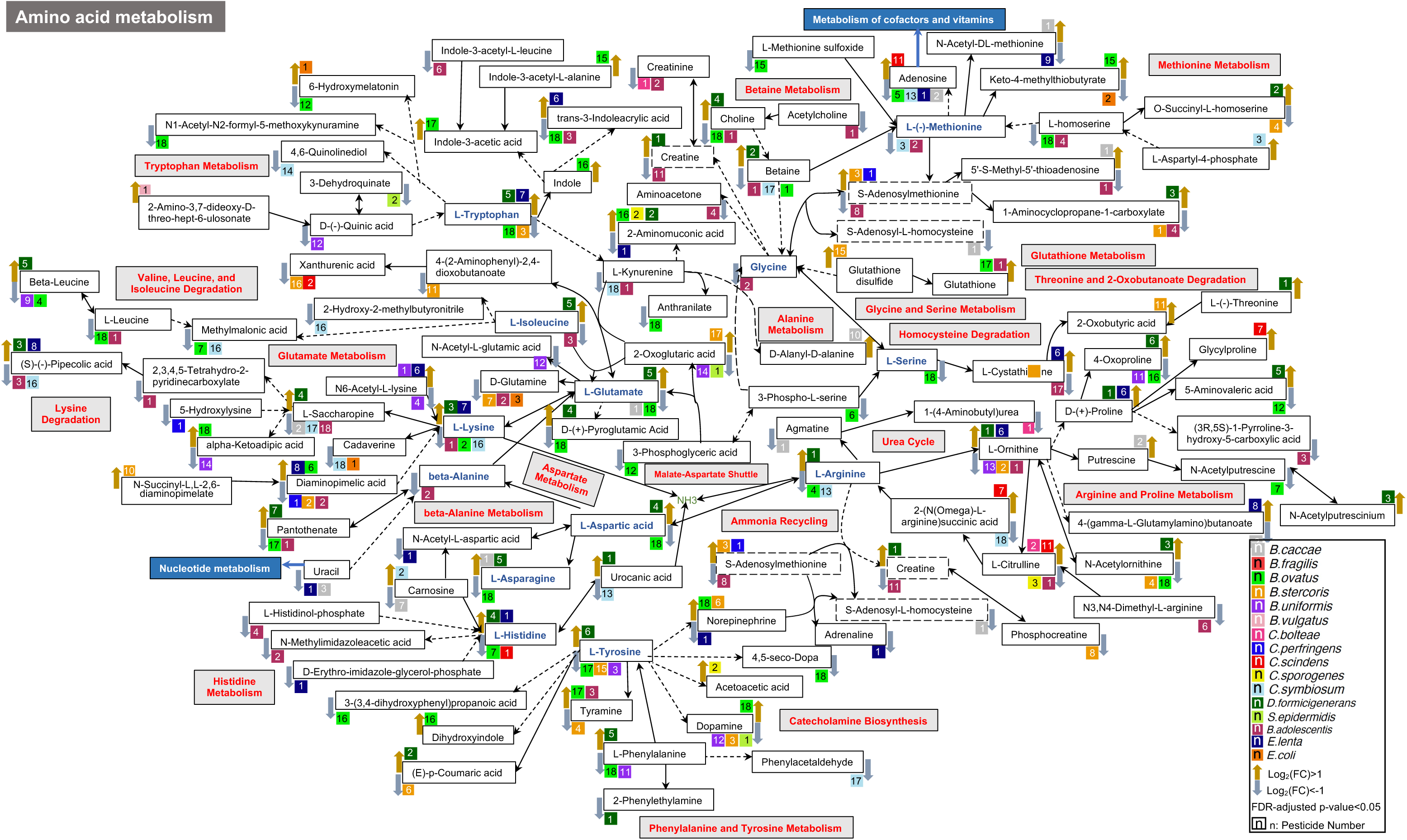
**pesticide-induced changes in amino acid metabolism related polar metabolites in gut microbiota.**

**Extended Fig.4:**
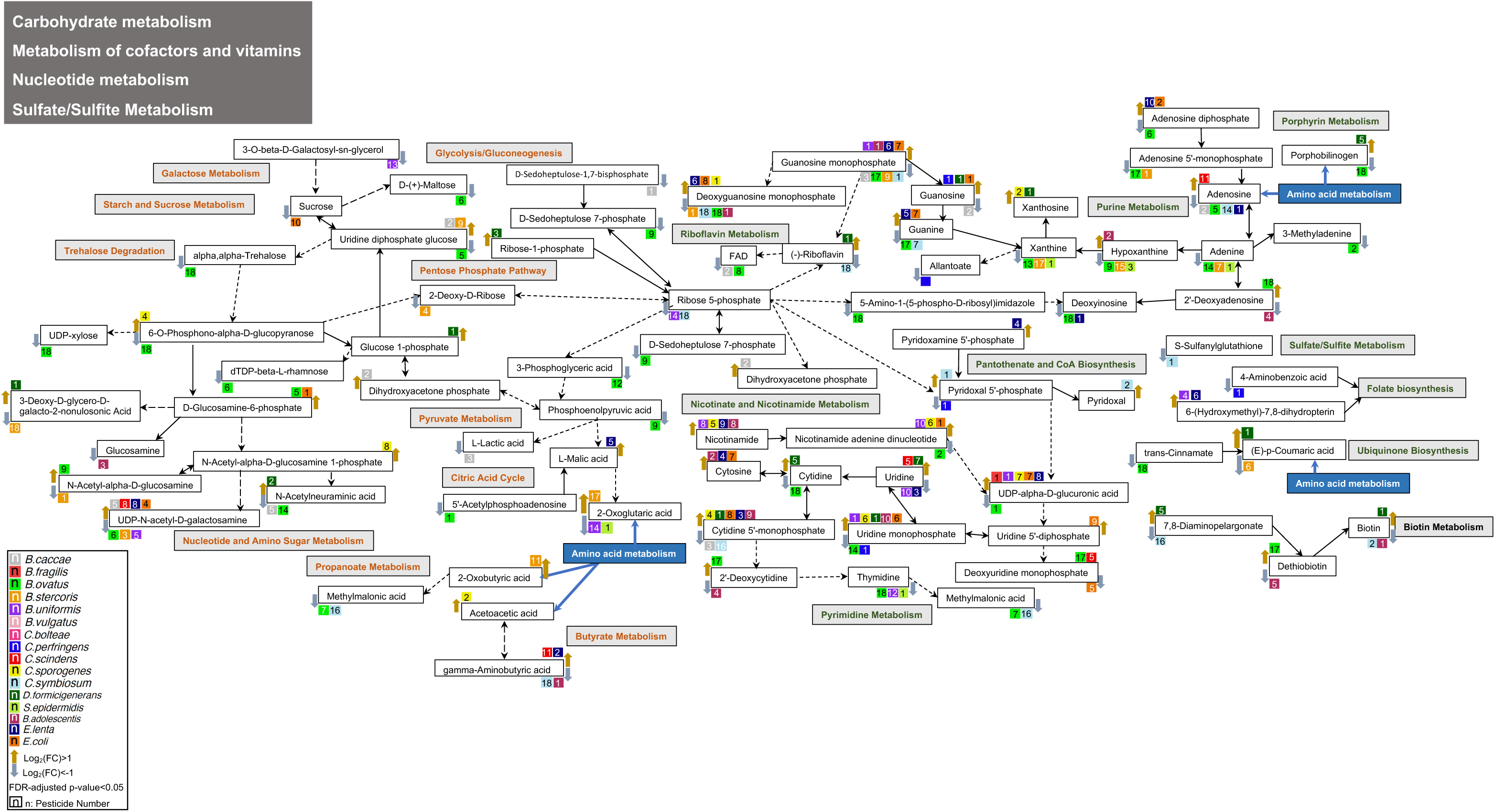
**pesticide-induced changes in carbohydrate metabolism, cofactor and vitamins metabolism, nucleotide metabolism and sulfate/sulfite metabolism related polar metabolites in gut microbiota.**

**Extended Fig.5:**
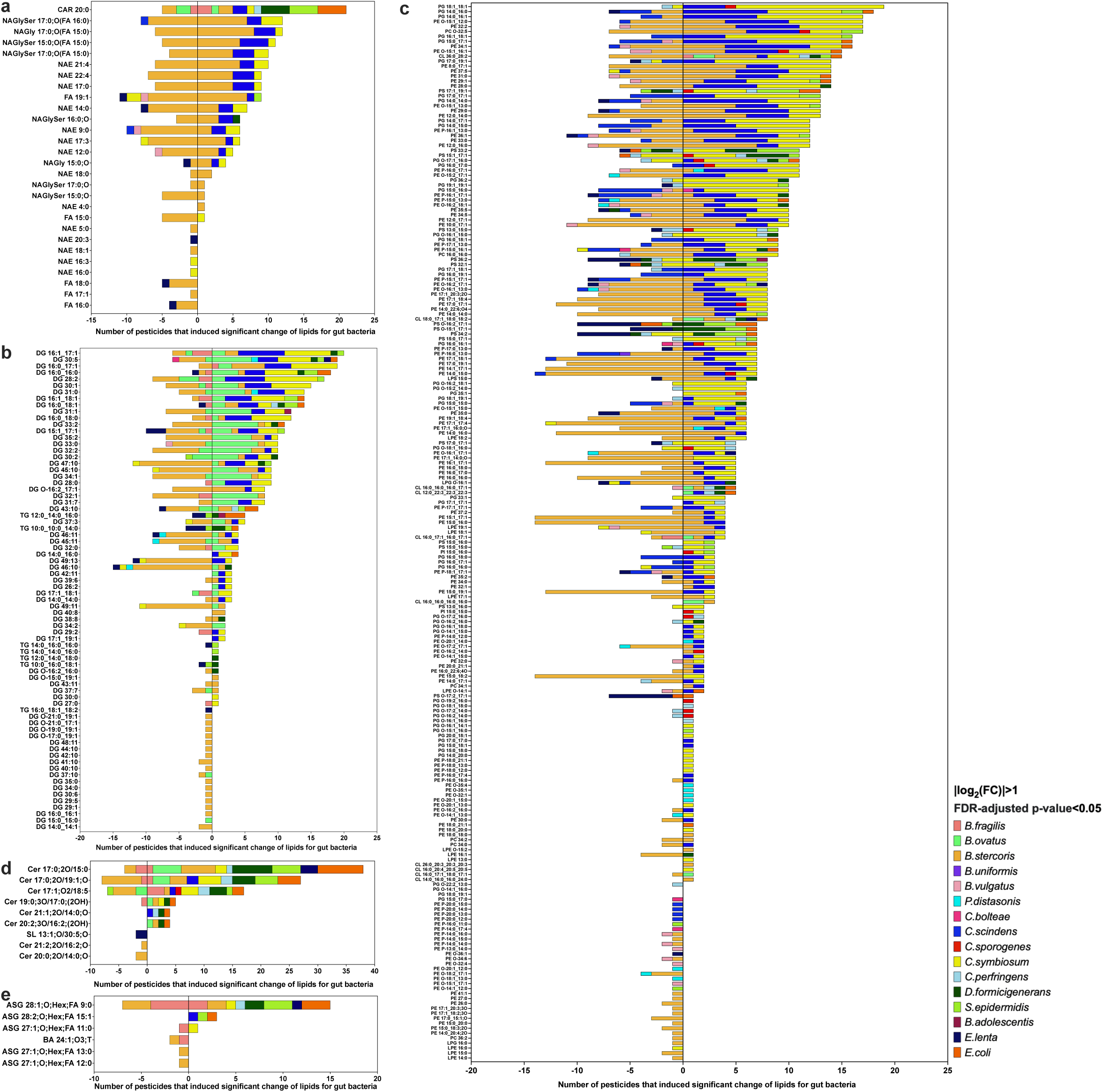
pesticide-induced changes in lipid metabolites in gut microbiota. a. pesticide-induced lipid changes for FA category in gut microbiota b. pesticide-induced lipid changes for GL category in gut microbiota c. pesticide-induced lipid changes for GP category in gut microbiota d. pesticide-induced lipid changes for SP category in gut microbiota e. pesticide-induced lipid changes for ST category in gut microbiota Data are presented as |log2(FC)|>1 and FDR-adjusted p-value <0.05 (a-d), (n=4). AHexBRS, Acylhexosyl brassicasterol; AHexCAS, Acylhexosyl campesterol; AHexCS, Acylhexosyl cholesterol; AHexSTS, Acylhexosyl stigmasterol; CAR, Acylcarnitine; Cer, ceramide; Cer_AP, Ceramide alpha-hydroxy fatty acid-phytospingosine; Cer_HS, Ceramide hydroxy fatty acid-sphingosine; Cer_HDS, Ceramide hydroxy fatty acid-dihydrosphingosine; Cer_NDS, Ceramide non-hydroxyfatty acid-dihydrosphingosine; Cer_NS, Ceramide non-hydroxyfatty acid-sphingosine; CL, cardiplipin; DG, diacylglycerol; DGDG, Digalactosyldiacylglycerol; DHSph, Sphinganine; EtherDG, Ether-linked diacylglycerol; EtherLPE, Ether-linked lysophosphatidylethanolamine; EtherLPG, Ether-linked lysophosphatidylglycerol; EtherPC, Ether-linked phosphatidylcholine; EtherPE Ether-linked phosphatidylethanolamine; EtherPG, Ether-linked phosphatidylglycerol; EtherPI, Ether-linked phosphatidylinositol; EtherPS, Ether-linked phosphatidylserine; FA, fatty acyl or fatty acid; FC, fold change; FDR, false discovery rate; HexCer, hexosylceramide alpha-hydroxy fatty acid-dihydrosphingosine; MGDG, Monogalactosyldiacylglycerol; GL, Glycerolipid; GP, Glycerophospholipid; LPE, ether-linked lyspphosphatidylethanolamine; LPG, Lysophosphatidylglycerol; LPS, Lysophosphatidylserine; NAE, N-acyl ethanolamines; NAGly, N-acyl glycine; NAGlySer, N-acyl glycyl serine; NAOrn, N-acyl ornithine; NATau, N-acyl taurine; oxFA, Oxidized fatty acid; oxTG, Oxidized triglyceride; oxPS, Oxidized phosphatidylserine; PC, phosphatidylcholine; PE, phosphatidylethanolamine; PEtOH, Phosphatidylethanol; PhytoSph, Phytosphingosine; PMeOH, Phosphatidylmethanol; PI, Phosphatidylinositol; PG, phosphatidylglycerol; PS, phosphatidylserine; SL, sulfonolipid; SM, Sphingomyelin; SP, Sphingolipid; SSulfate, Sterol sulfate; ST, sterol lipid; TG, triacylglycerol.

**Extended Fig.6:**
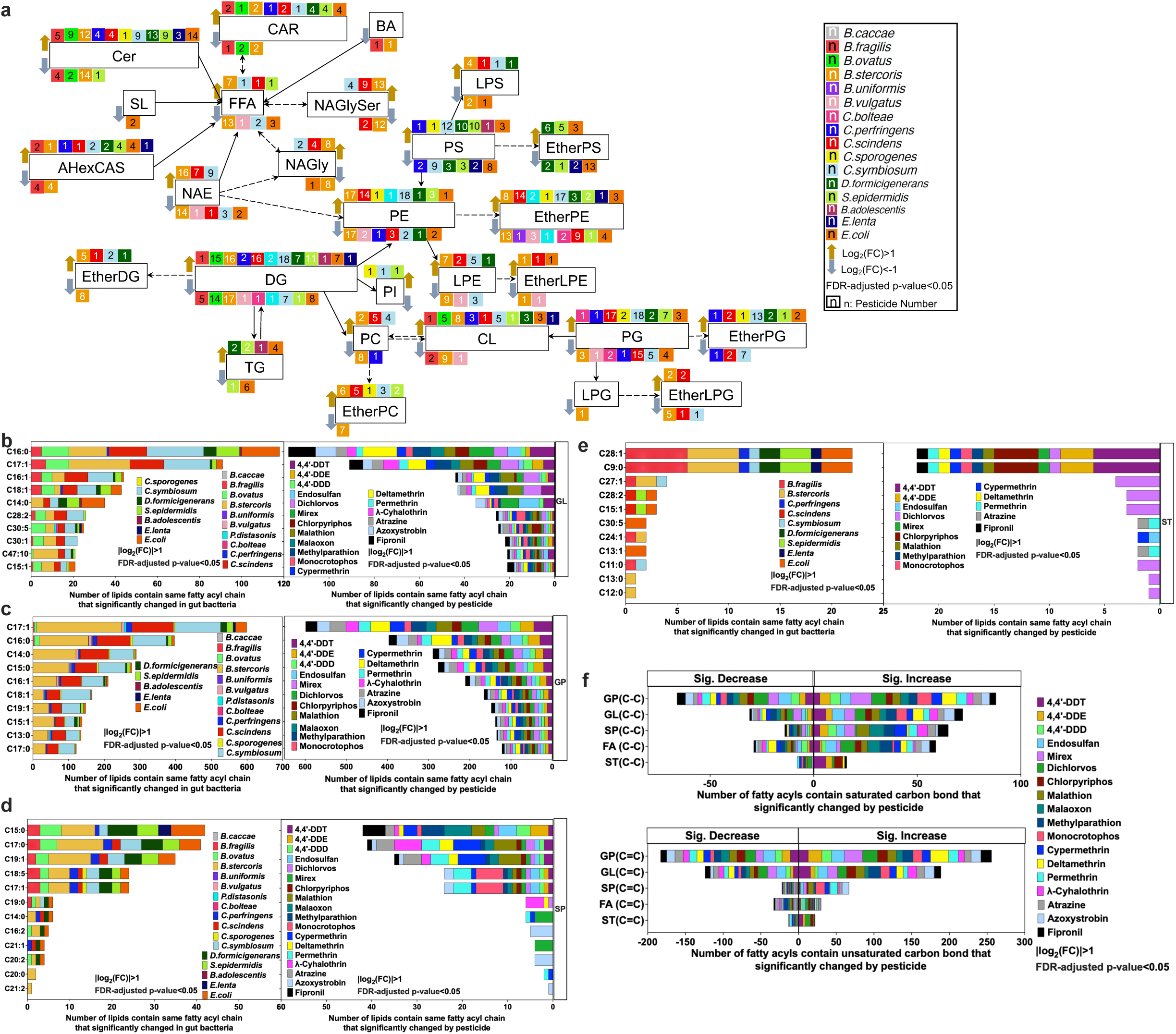
pesticide-induced changes in lipid metabolites in gut microbiota. a. Changes in the primary lipid classes participate pathways after pesticide exposure on gut bacteria strains. b-e. Number of lipids contain same fatty acyl chain that significantly changed for GL, GP, SP, ST category. f. Number of lipid carbon bond changes for each pesticide in gut bacteria Data are presented as |log2(FC)|>1 and FDR-adjusted p-value <0.05 (a-d) (n=4). CAR, Acylcarnitine; Cer, ceramide; CL, cardiplipin; DG, diacylglycerol; DGDG, Digalactosyldiacylglycerol; DHSph, Sphinganine; EtherDG, Ether-linked diacylglycerol; EtherLPE, Ether-linked lysophosphatidylethanolamine; EtherLPG, Ether-linked lysophosphatidylglycerol; EtherPC, Ether-linked phosphatidylcholine; EtherPE Ether-linked phosphatidylethanolamine; EtherPG, Ether-linked phosphatidylglycerol; EtherPI, Ether-linked phosphatidylinositol; EtherPS, Ether-linked phosphatidylserine; FA, fatty acyl or fatty acid; FC, fold change; FDR, false discovery rate; GL, Glycerolipid; GP, Glycerophospholipid; LPE, ether-linked lyspphosphatidylethanolamine; LPG, Lysophosphatidylglycerol; LPS, Lysophosphatidylserine; NAE, N-acyl ethanolamines; PC, phosphatidylcholine; PE, phosphatidylethanolamine; PI, Phosphatidylinositol; PG, phosphatidylglycerol; PS, phosphatidylserine; SL, sulfonolipid; SM, Sphingomyelin; SP, Sphingolipid; ST, sterol lipid; TG, triacylglycerol.

**Extended Fig.7.**
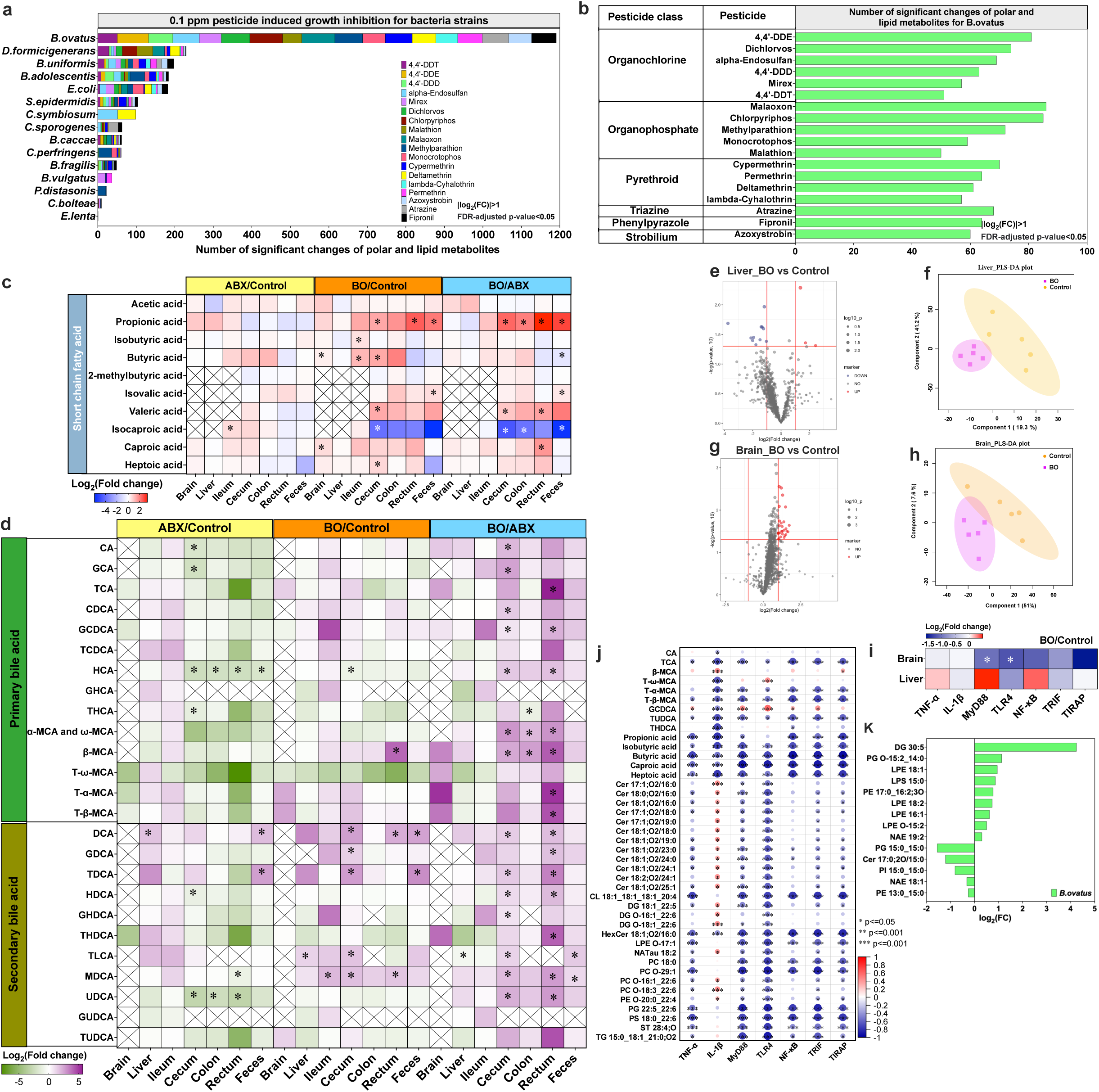
4,4’-DDE induced metabolic changes in *B.ovatus* transplanted C57BL/6 mice. a, number of significant changes in polar and lipid metabolites in the gut bacteria strains inhibited by pesticides at 0.1 μg/mL. b, number of significant changes in polar and lipid metabolites in B.ovatus after pesticides exposure. c, targeted analysis of short-chain fatty acid levels in organs and tissues of mice at the end of experiment in all groups. d, targeted analysis of bile acid levels in organs and tissues of mice at the end of experiment in all groups. e, volcano plot of significant changes in polar and lipid metabolites in liver between the BO and Control group (|log2(FC)|>1 and p-value <0.05). f, PLS-DA separation of liver between the BO and Control group. g, volcano plot of significant changes in polar and lipid metabolites in brain between the BO and the Control group (|log2(FC)|>1 and p-value <0.05). h, PLS-DA separation of brain between the BO and Control group. i, mRNA relative expression of receptors in signaling pathways in the brain and liver of C57BL/6 mice at the end of experiment. j, Pearson correlation analysis between lipids and receptors in the brain at the end of experiment. k, Significant changes of lipids for *B.ovatus* after 4,4’-DDE exposure. Data (a-b) are presented as |log2(FC)|>1 and FDR-adjusted p-value <0.05 (a-d). Data (c) are presented as mean ± SEM (n=5). p values were calculated by t-test, and p <0.05 (*) represents statistically significant. FC, fold change; FDR, false discovery rate. PLS-DA, partial least squares-discriminant analysis.

## Data reporting

No statistical methods were used to predetermine sample size. The experiments were not randomized, and the investigators were not blinded to allocation during experiments and outcome assessment.

## Growth conditions

The bacteria utilized in this study were obtained from the American Type Culture Collection (ATCC) and Coli Genetic Stock Center *(K12)* grown in Gifu anaerobic medium (GAM broth) in Coy anaerobic. The bacterial strains employed in this research can be found in **Table S1**. Anaerobic conditions were maintained using a nitrogen and hydrogen gas mixture, ensuring oxygen levels between 0 and 20 parts per million (ppm), alongside 2-3% hydrogen. The anaerobic state was monitored using the anaerobic monitor CAM-12. To eliminate oxygen and produce water molecules, two Stak-Pak systems equipped with palladium catalysts were employed. A vacuum airlock was utilized to minimize oxygen levels when transferring reagents or materials into and out of the glove box. Additionally, surfaces of equipment and materials were disinfected using 10% bleach and 75% alcohol. Glycerol stock of all bacterial strains was revived for plate streaking, and a single colony was transferred to GAM broth. Experimental cultures were initiated from the second passage culture following inoculation.

## Pesticide dose-dependent growth inhibition

To evaluate the growth inhibition of individual bacteria species under anaerobic conditions, a concentration gradient (0.05 μg/mL, 0.1 μg/mL, 0.5 μg/mL, and 1 μg/mL) was employed for all pesticides. For most of tested pesticides, the 0.05-1 μg/mL concentrations were chosen to ensure it remained within the acceptable limits set by the pesticide MRLs database from the US, EU and China, and represented a potential exposure concentration in the gastrointestinal tract (**Extended Fig. 1a**). Triplicate screenings for each bacterium-pesticide interaction were conducted within a 96-well deep plate. Prior to pesticide exposure, fresh Gifu anaerobic medium (GAM broth) was introduced into the anaerobic chamber the evening before. In order to assess dose-dependent growth inhibition caused by the pesticides, a 5 μL work solution of the pesticide, diluted with dimethyl sulfoxide (DMSO), or 5 μL of DMSO as a control, along with 1 mL of a second passage bacteria culture to achieve a starting optical density (OD) of 0.05 at 590 nm, were added to the 96- well deep plates under anaerobic conditions. The mixture was gently pipetted until thoroughly mixed and incubated at 37°C for 3-5 hours, depending on the growth rate of the bacteria. The growth of the bacteria was monitored by measuring the OD at 590 nm using a BioTek cytation 5. Statistical significance was assessed using a T-test with a p-value cut-off of 0.05.

## Pesticide bioaccumulation detection and metabolism evaluation

Single bacteria strain from second passage culture were inoculated at a starting OD590 nm of 0.05 of 1 ml GAM culture containing 0.1 μg/mL pesticide in 96 well plates and incubated for 12 h while shaking at 37 °C under anaerobic condition. Plates were closed with lids.

To detect bioaccumulation, a combinatorial pooling strategy ^73^ was employed to allocate 18 pesticides into 8 pools, ensuring that each pesticide was represented in quadruplicate (**Extended Fig. 2a**). After 12 hours, the 96-well plates containing the bacteria culture were sealed and removed from the anaerobic chamber for storage at -80°C until analysis. To determine the pesticide concentration in the bacterial strains using GC-QQQ MS, the 96-well plate needed to be thawed at 4°C for 15 hours, followed by centrifugation at 4400 rpm for 20 minutes. The supernatant was discarded, and the pellet was washed three times with 1 mL of PBS (pH 7.4). A pesticide extraction solution was prepared using acetonitrile and 0.050 μg/mL of chlorpyriphos-methyl as an internal standard. After adding 300 μL of the extraction solution and vortexing for 1 minute, 300 μL of the mixture was transferred into a 2 mL Eppendorf tube containing 0.5 g of NaCl. The same steps were repeated by adding 300 μL of the extraction solution back into the 96-well plate. Finally, the 600 μL of the extraction solution was combined and vortexed for 3 minutes. After centrifugation at 10,000 r/min for 3 minutes, all the supernatant was transferred into a 2 mL dSPE tube with two 3mm glass beads for purification. The sample was homogenized for 15 seconds using a Beadbeater (MiniBeater-16, Model 507) for a total of three cycles, followed by centrifugation at 10,000 r/min for 3 minutes. The resulting supernatant was transferred into 2 mL glass vials with glass inserts for GC-QQQ MS analysis.

To assess metabolism, 18 individual pesticides were added in quadruplicate to a 96-well plate, along with 1 mL of a single bacteria culture at a starting OD590nm of 0.05. Two plates with the same pesticide combinations were prepared for subsequent metabolomics and lipidomics analyses. After 12 hours, the 96-well plates were covered with sealing films and removed from the anaerobic chamber, then stored at -80°C until analysis. Prior to extraction, the frozen 96-well plates were thawed at 4°C for 15 hours and subsequently centrifuged at 4400 rpm for 20 minutes. The supernatant was discarded, and the pellet was washed three times with 0.5 mL of PBS (pH 7.4). For metabolomics extraction, an extraction solution containing methanol and 200 μg/mL of 13C,15N-amino acids as an internal standard were prepared. After adding 300 μL of the extraction solution to the 96-well plates, the samples were vortexed for 3 minutes and kept at -20°C for 20 minutes. Following centrifugation at 4400 rpm and 4°C for 20 minutes, the supernatant was collected in a 2 mL glass vial with an insert for UPLC-QE orbitrap analysis. As for lipidomics extraction, the extraction solution was a mixture of 2-propanol and 0.275 μg/mL of 13C-labeled lipids as an internal standard. After adding 290 μL of the lipid extraction solution, the samples were vortexed for 2 minutes and ultrasonicated in ice water for 20 minutes, followed by standing at 4°C for 30 minutes. After centrifugation at 4400 rpm and 4°C for 20 minutes, the supernatant was collected in a 2 mL glass vial with an insert for UPLC-QE orbitrap MS analysis.

## Mouse experiment

All animal experiments conducted in this study were carried out following the approved protocols by the Ohio State University Institutional Laboratory Animal Care and Use Committee. Male C57BL/6 mice (n = 32) aged 7 weeks and obtained from Jackson Lab were housed in a controlled environment at 25°C with a 12-hour light-dark cycle. Following a one-week acclimation period, the mice were randomly assigned to three groups, with 5 mice per group. The mice were provided with sterile food and water ad libitum throughout the study. The four groups were designated as ABX, Control, and BO, corresponding to specific periods as outlined below.

**Period 1**: a pseudo germ-free mouse model was established by treating all groups of mice with broad-spectrum antibiotics. At 8 and 9 weeks of age, the mice received a continuous administration of ampicillin (1 g/L), neomycin sulfate (1 g/L), and metronidazole (1 g/L) obtained from Sigma-Aldrich Co. Ltd, USA, which were dissolved in their drinking water. This antibiotic treatment lasted for 14 consecutive days. The drinking solution was replaced and documented every 2 days, while sterile food was refreshed and recorded on a weekly basis.

**Period 2**: In the mono-colonization experiments, B. ovatus was cultured in an anaerobic chamber and their identities were confirmed through phenotype verification using GAM plate streaking and PCR primers. For B. ovatus, the primers used were as follows: Forward: 5’-3’ TGCAAACTRAAGATGGC and Reverse: 5’-3’ CAAACTAATGGAACGCATC. After culturing for 24 hours, a 10 mL saturated bacterial culture was centrifuged and washed three times with PBS. The bacterial pellets were then resuspended in 10 × 1 mL sterile PBS in 2 mL sterile Eppendorf tubes. In the BO group, mice were colonized with *B. ovatus* ATCC8483 (approximately 8 × 10˄6 colony-forming units (CFU)) obtained from the saturated cultures. This colonization process involved orally administering 200 μL of the respective cultures through gavage. On the other hand, mice in the ABX and Control groups were only orally gavaged with 200 μL of PBS. Fecal samples were collected both before and after mono-colonization for phenotype verification. The sterile water was replaced and documented every 2 days, and sterile food was refreshed and recorded on a weekly basis.

**Period 3**: Regarding pesticide exposure, once successful mono-colonization was achieved, the mice from the Control, and BO groups were subjected to a 4-week exposure to 0.1 μg/mL of 4,4’- DDE in their drinking water. Meanwhile, the mice in the ABX group received sterile water without pesticide. The drinking water was replaced and documented every 2 days, and sterile food was refreshed and recorded on a weekly basis. Prior to and after mono-colonization, 16S amplicon sequencing was performed on the V4 region (515F, 806R) of microbial populations found in individual mice’s feces. Following the completion of the above-mentioned three periods, the mice were euthanized using CO2. Thirteen different sample types (serum, brain, liver, lung, heart, spleen, kidney, testis, ileum, cecum, colon, rectum, and feces) were collected and weighed from each mouse. All samples were divided into four replicates and stored at -80°C until further analysis. In the mouse experiments, a single biological replicate represents a specific sample type, such as serum, obtained from an individual mouse. This means that each biological replicate is derived from a different mouse. Prior to euthanization, feces were collected, and blood samples were taken from the heart and centrifuged to obtain serum for subsequent analysis. The contents from the ileum, cecum, colon, and rectum were thoroughly removed.

## Mouse samples preparation for pesticide analysis

To begin the extraction process, take 50 μl of serum, 100-300 mg of feces, or 10-400 mg of tissues and transfer them into a 2 mL dSPE tube containing 2 beads for pesticide extraction. Add 800- 1000 μl of acetonitrile to the tube, adjusting the volume based on the mass or volume of the samples. Vortex the mixture using a beadbeater for 15 seconds (or 60 seconds for samples from the ileum, cecum, colon, or rectum), repeating this step three times. Subsequently, centrifuge the tube at room temperature and 12,000 rpm for 10 minutes. Carefully transfer the resulting supernatant, which should be around 600-800 μL, into 2 mL Eppendorf tubes for further processing using a speedvac. To redissolve the sample, add 100 μl of isooctane and the internal standard (chlorphriphos-methyl at a concentration of 0.05 μg/mL). Centrifuge the mixture once again at room temperature and 12,000 rpm for 10 minutes. Transfer the supernatant into a 2 mL glass vial with a glass insert for subsequent analysis. Finally, subject the samples to GC-QQQ MS to detect 4,4’-DDE and its metabolites 4,4’-DDD.

## Mouse samples preparation for SCFAs, BAs, and metabolomics analysis

The process involved weighing 50 μL of serum, 100-200 mg of feces, and 10-100 mg of tissue samples into 2 mL polypropylene microvials. Two glass beads and 300-500 μL of methanol with 200 μg/mL of 13C,15N-amino acids were added to facilitate homogenization using a beadbeater for 15 seconds (repeated three times with 60-second intervals). The mixture was then sonicated in ice water for 30 minutes and left at -20°C for 20 minutes. Following centrifugation at 14,000 rpm and 4°C for 20 minutes, 180 μL of supernatant was transferred into a 2 mL glass vial with an insert for bile acids (BAs) and metabolomics analysis. For SCFAs (short-chain fatty acids) derivatization, 40 μL of supernatant from serum, feces, liver, brain, ileum, cecum, colon, and rectum was transferred into a 2 mL Eppendorf tube. This step followed our previously published method ^74^. To summarize, either a 40 μL standard solution or a 40 μL supernatant was thoroughly mixed with 20 μL of a 200 mM 3NPH·HCl solution and 20 μL of a 120 mM EDC·HCl-6% pyridine solution. The samples were vortexed for 2 minutes and then incubated in a 40°C water bath for 30 minutes. Following that, the samples were cooled on ice for 1 minute. Depending on the sample, either 420 μL or 920 μL of a 10% acetonitrile solution was used to dilute the derivatized samples before conducting UPLC-QE Orbitrap MS analysis. Due to the high concentration of acetic acid in the samples, all diluted samples still required a further dilution of 20 times before reanalysis by UPLC- QE Orbitrap MS.

## Mouse samples preparation for lipidomics analysis

To begin, approximately 20 mg samples of kidney, lung, heart, testis, ileum, cecum, colon, rectum, and feces, along with 20 μL of serum, were weighed into 2 mL microvials. A lipid extraction solution consisting of 200 μL of 2-propanol with 0.275 μg/mL 13C-labeled lipids was added to the microvials. For brain samples weighing 10-100 mg and liver samples weighing 75-350 mg, 300 μL and 1 mL of the lipid extraction solution, respectively, were added to facilitate sample extraction. In all vials, 2 glass beads were included for homogenization using a Beadbeater, either for 15 seconds or 60 seconds, repeated three times. The homogenized samples were then subjected to ultrasonication in ice water for 20 minutes and left to stand overnight at -20°C. The following day, the samples were centrifuged at 14,000 rpm and 4°C for 10 minutes, and the resulting supernatant was collected into a 2 mL glass vial with an insert for UPLC-QE Orbitrap MS analysis.

### GC-QQQ MS methods

Data acquisition was performed using an Agilent 8890 GC system coupled with a 7010 Triple Quadrupole mass spectrometer (GC-QQQ MS), which was equipped with a Gerstel autosampler. Data analysis was conducted using Agilent MassHunter Quantitative analysis software. For separation, a J&W DB-5MS column with a 95% dimethyl/5% diphenyl polysiloxane composition, measuring 30 meters in length, 0.25 millimeters in internal diameter, and coated with a 0.25- micrometer film, was used. Additionally, a 10-meter empty DuraGuard guard column was employed. Samples of 1 μL were injected using an Agilent ultra-inert inlet liner with a 4 mm ID, utilizing a splitless, single taper, wool configuration. The inlet temperature was set to 250°C with splitless mode, and the gas saver was set to 20 mL/min after 3 minutes. Helium gas was used as the carrier gas with a flow rate of 1.005 mL/min. Nitrogen served as the collision gas at a rate of 1.5 mL/min, while helium was utilized as the quenching gas at 4 mL/min. The initial oven temperature was 60°C for 1 minute, followed by an increase of 40°C/min to 170°C, and then an increase of 10°C/min to 310°C, which was held for 10 minutes. The total run time was 27.75 minutes. For the post-run period, the temperature was set at 320°C, and the flow rate was maintained at 1.2 mL/min for 5 minutes. The MSD transfer line temperature was 280°C, the ion source temperature was 280°C, the quadrupole temperature was 150°C, and a gain factor of 1 was applied. In dMRM (dynamic multiple reaction monitoring) mode, 18 pesticides, each with at least two pairs of precursor and product ions, were selected for both qualitative and quantitative analysis. The mass resolution of both MS1 and MS2 was set to wide. A targeted pesticide method was established and validated. As an initial step in evaluating the analytical performance, the pesticide method was verified to exhibit good linearity and recovery, as demonstrated in **Extended Fig. 2c**. To determine the final pesticide concentrations in samples obtained from in vivo or in vitro experiments, a 50 μg/mL Chlorpyriphos-methyl internal standard was utilized for calculation purposes.

### UPLC-QE Orbitrap MS methods

This study utilized a Thermo Vanquish UPLC system coupled with a Q-Exactive Orbitrap mass spectrometer (UPLC-QE Orbitrap MS) equipped with a heated electrospray ionization (HESI) probe from Thermo Fisher in California, USA. For metabolomics analysis, a Waters Xbridge BEH Amide 2.5μm 2.1×150mm column was employed to separate polar metabolites in both negative and positive ionization modes, with separate injections for each mode. Mobile phase A consisted of a mixture of acetonitrile and water (10/90, v/v), containing 5 mM ammonium acetate and 0.1% acetic acid. Mobile phase B comprised a mixture of acetonitrile and water (90/10, v/v), also containing 5 mM ammonium acetate and 0.1% acetic acid. A linear gradient elution program was employed, starting with 70% B from 0 to 0.1 minutes, decreasing to 30% B from 0.1 to 5 minutes, holding for 4 minutes, and then returning to 70% B by 2 minutes and holding for 9 minutes. The total run time was 20 minutes, with a flow rate of 0.30 mL/min and a column temperature of 40°C. The data collection resolution for full scan analysis was set to 70,000 within the m/z range of 60- 900. The automatic gain control (AGC) target was 3e6, and the maximum ion trap (IT) time was 200 ms.

For bile acids (BAs) analysis, a Phenomenex Kinetix C18 column (2.6 µm, 150 mm × 4.6 mm ID) was used for separation in negative ionization mode, following our previously reported method ^75^. The mobile phase A consisted of a methanol:acetonitrile:water mixture (1:1:3, v/v/v) with 1 mM ammonium acetate and 0.1% acetic acid, while mobile phase B consisted of a methanol:acetonitrile:2-propanol mixture (4.5:4.5:1, v/v/v) with 0.1% acetic acid. Gradient elution was applied, and MS1 and MS2 data were collected using the PRM mode and t-SIM mode.

For SCFAs analysis, CSH C18 1.7μm 2.1×100mm column (Waters Corp, Milford, MA, USA) was applied for SCFAs separation in negative ionization mode ^74^. The mobile phase A consisted of water with 0.1% formic acid, and mobile phase B comprised acetonitrile with 0.1% formic acid. MS1 and MS2 data were acquired using the PRM mode and t-SIM mode.

For lipidomic analysis, an Acquity UPLC CSH C18 1.7μm 2.1×100mm column (Waters Corp, Milford, MA, USA) was utilized for separation of lipid metabolites in both negative and positive ionization modes, following our previously reported method ^76^. Mobile phase A consisted of a mixture of acetonitrile and water (60/40, v/v), containing 10 mM ammonium acetate and 0.1% formic acid. Mobile phase B comprised a mixture of acetonitrile and 2-propanol (10/90, v/v), also containing 10 mM ammonium acetate and 0.1% formic acid. Full scan/ddMS^2^ mode was used to acquire MS1 and MS2 information.

### Transcriptomics analysis

Liver and brain samples were homogenized in TRIzol reagent at a ratio of 1 mL per 50-100 mg of tissue. Following homogenization, the mixtures were incubated at room temperature for 5 minutes. Next, 100 μL of BCP was added to the mixture and incubated again at room temperature for 5 minutes. The samples were then centrifuged at 14,000 rpm and 4 °C for 8 minutes, and the resulting supernatant was carefully transferred to a fresh 1.5 mL microtube. To precipitate the RNA, 250 μL of a high salt precipitation solution and 250 μL of 2-propanol were added and incubated at room temperature for 10 minutes. After centrifugation at 14,000 rpm and 4 °C for 5 minutes, the supernatant was aspirated off, and the RNA pellets were air-dried at room temperature for approximately 5-10 minutes. Finally, 50-200 μL of DEPC water was used to dissolve the RNA. After quantifying the RNA, the concentration was adjusted to 200 ng/μL for cDNA synthesis using the iScript cDNA synthesis kit from Bio-Rad. The synthesized cDNA was stored at -20 °C for qRT-PCR analysis. 10 primer pairs were compiled and listed in **Table S9**.

### 16s RNA sequencing

Briefly, PCR amplicon libraries targeting the 16S rRNA encoding gene present in metagenomic DNA are produced using a barcoded primer set adapted for the Illumina HiSeq2000 and MiSeq. Specifically, the V4 region of the 16S rRNA gene (515F-806R) is PCR amplified with region-specific primers that include sequencer adapter sequences used in the Illumina flowcell. Each 25 µL PCR reaction contains 9.5 µL of MO BIO PCR Water (Certified DNA-Free), 12.5 µL of QuantaBio’s AccuStart II PCR ToughMix (2x concentration, 1x final), 1 µL Golay barcode tagged Forward Primer (5 µM concentration, 200 pM final), 1 µL Reverse Primer (5 µM concentration, 200 pM final), and 1 µL of template DNA. The conditions for PCR are as follows: 94 °C for 3 minutes to denature the DNA, with 35 cycles at 94 °C for 45 s, 50 °C for 60 s, and 72 °C for 90 s; with a final extension of 10 min at 72 °C to ensure complete amplification. Amplicons are then quantified using PicoGreen (Invitrogen) and a plate reader (Infinite® 200 PRO, Tecan). Once quantified, volumes of each of the products are pooled into a single tube so that each amplicon is represented in equimolar amounts. This pool is then cleaned up using AMPure XP Beads (Beckman Coulter), and then quantified using a fluorometer (Qubit, Invitrogen). After quantification, the molarity of the pool is determined and diluted down to 2 nM, denatured, and then diluted to a final concentration of 6.75 pM with a 10% PhiX spike for sequencing on the Illumina MiSeq. Amplicons are sequenced on a 151bp x 12bp x 151bp MiSeq run using customized sequencing primers and procedures.

## Data analysis

### Masshunter quantitative analysis

Data analysis of GC-QQQ MS was conducted using Agilent MassHunter Quatitative Analysis version 10.2 software (Agilent Technologies). A new analysis method was generated based on the acquired MRM data. The software automatically populated the precise precursor and product ions as well as the retention time in certain sections of the data analysis method. The quantitative ion pairs and retention time for both in vitro (15 pesticides and 3 metabolites) and in vivio (4,4’-DDE and 4,4’-DDD) samples were manually verified and adjusted if necessary. Subsequently, the method was saved and utilized for batch quantification of all samples.

### Compound discover analysis

For the metabolomics analysis, the Compound Discover software (version 3.3, Thermo Fisher Scientific) was employed to process and analyze the ’.raw’ data files obtained from the UPLC-QE Orbitrap MS. To initiate the data processing task, a new study was created, and a customized non-targeted metabolomics workflow was employed for data processing. The ’.raw’ data files were added to the project and categorized into three predefined groups: Blank, Samples, and Identification only. The identification of compounds was performed using mzCloud (ddMS2), ChemSpider (formula or exact mass), and an in-house database containing m/z values of 171 standards and 20 13C15N amino acids. The workflow included retention time correction, feature detection, and chromatogram alignment. The parameters used were as follows: polarity (positive [M+H]+ or negative [M-H]-) determined by the raw data, maximum shift of 0.2 min, mass tolerance of 5 ppm, and a minimal peak intensity of 1 × 104. After data processing, a characteristic table was generated based on the m/z and retention time of each molecule, which provided the peak areas of each compound across all samples. Subsequently, the data was normalized and exported in.csv format. Quality control (QC) samples were utilized to calculate the coefficient of variation (CV) for each compound, and compounds with CV values less than 20% were selected for further statistical analysis.

### MS-DIAL analysis

The lipidomics analysis involved the utilization of the MS-DIAL software (version) to analyze all in vitro and in vivo data acquired from the UPLC-QE Orbitrap MS. The raw data acquisition was performed using Xcalibur 4.0 software (Thermo Fisher Scientific, USA). Subsequently, the raw data from the DDA and DIA experiments were converted from the vendor-specific file format (.raw) to the Analysis Base File format (.abf) using the freely available Reifycs ABF converter (https://www.reifycs.com/AbfConverter/).

After conversion, the MS-DIAL software (version 4.24) was employed for various data processing tasks, including feature detection, spectral deconvolution, peak identification, and alignment between samples. Quality control samples (QC) from each bacteria strain or mice samples were used for peak alignment. During the analysis, specific adducts were selected based on the ionization mode. In positive ionization mode, adducts such as [M+H]+, [M+NH4]+, [M+Na]+, [M+ACN+H]+, [M+H-H2O]+, [M+H-2H2O]+, [2M+H]+, and [M+2H]2+ were chosen, while in negative ionization mode, adducts including [M-H]-, [M-H2O-H]-, [M+Na-2H]2-, [M+FA-H]-, [M+Hac-H]-, [2M-H]-, and [M-2H]2- were selected. The lipid database settings were kept as default for both positive and negative ion modes.

Chemical assignment of molecular features in the samples was performed by comparing the recorded retention time (RT) and m/z information to the reference library constructed from authentic standards. Tolerance windows of 0.05 min for RT and 0.01 Da for m/z were set. To filter the results, a minimal peak count filter of 5,000 or a signal-to-noise ratio (S/N) filter of 10 was applied to all samples. The MS-DIAL analysis generated a comprehensive list of metabolite names, m/z values, RT values, formulas, ontologies, INCHIKEYs, SMILES representations, S/N ratios, and peak areas for high confidence annotations, as well as all unknown molecular features for both positive and negative polarity modes. Specific metabolite features were excluded from the list under the following conditions: (1) if they were detected only in the blank controls, (2) if the coefficient of variation (CV) in the QC samples was higher than 20%, (3) if the annotated compounds were identified in both positive and negative polarity modes and had lower peak areas or S/N ratios, or higher CV values, and (4) if the molecular features were unknown, they were also removed for further analysis.

### Statistics and reproducibility

The results were presented as mean ± SE, statistical analysis and data visualization were conducted using GraphPad Prism 8 software, OriginLab 2020, Biorender, and R Studio. For normally distributed data, a two-tailed unpaired Student’s t-test was performed. To control the false discovery rate (FDR), the p-values were adjusted using the Benjamini-Hochberg (BH) method. A resulting adjusted p-value of less than 0.05 was considered statistically significant. TidyMass software was utilized for the analysis of metabolomics and lipidomics data ^77^. To conduct data analysis, three data frames were created: expression_data, sample_info, and variable_info. These data frames were utilized by the libraries (massdataset, massstat, metpath) to generate various plots such as PCA plots, volcano plots, pathway enrichment bars, and pathway enrichment scatter plots. Each experimental group was compared with its respective control group. After performing t-tests and FDR correction, metabolites with a p-value < 0.05, along with their adjusted p-values and fold change information, were exported as.csv files for further analysis. Pathway changes were evaluated using a significance threshold of p-value < 0.05. The pathway name, p-value, adjusted p-value, and mapped metabolite IDs were exported as CSV files using R code for further analysis. For metabolite analysis, metabolites and lipids were filtered based on an adjusted p-value < 0.05. Heatmaps were generated using log2(fold change) values for both in vitro and in vivo experiments. Given the complex composition of lipids, including different functional groups, fatty acyl chains, and double bonds, these characteristics were also summarized and depicted in the heatmap based on log2(fold change). For pathway analysis, pathways with an adjusted p-value < 0.05 were sorted and visualized in heatmaps for both in vivo and in vitro studies.

## Data availability

The raw metabolomics data for the in *vitro* and in *vivo* experiments can be accessed publicly on MassIve under study number MSV000095526, MSV000095539, and MSV00095671. All supplementary data related to this study has been compiled and can be found in **Table S1-S9**.

## Code availability

We developed custom R code to facilitate the processing and visualization of the UPLC-QE Orbitrap MS data obtained from the in vitro and in vivo experiments. All original code has been uploaded into https://data.mendeley.com/datasets/9f5xzypspc/1 (DOI:10.17632/9f5xzypspc.1) for peer review. This code provides a comprehensive toolkit for data analysis and facilitates the exploration of various aspects of the metabolomics and lipidomics datasets.

